# Disturbance increases diversity by weakening competition rather than reducing intrinsic population growth

**DOI:** 10.64898/2026.09.07.749828

**Authors:** Yuval Neumann, Or Gross, Tal Schabes, Niv DeMalach, Michael Kalyuzhny

**Affiliations:** Institute of Plant Sciences and Genetics in Agriculture, Robert H. Smith Faculty of Agriculture, Food and Environment, The Hebrew University of Jerusalem, Rehovot 7610001, Israel; Department of Ecology, Evolution and Behavior, Alexander Silberman Institute of Life Sciences, Faculty of Science, The Hebrew University of Jerusalem, Jerusalem 9190401, Israel

## Abstract

Physical disturbance is pervasive across ecosystems. However, why it can increase diversity remains unresolved. For over five decades, two hypotheses have highlighted contrasting mechanisms: weakened heterospecific competition, or reduced intrinsic demographic rates that slow competitive exclusion. Here, we formalize these largely verbal hypotheses using standard models of multispecies community dynamics and test the resulting quantitative predictions in an experimental mesocosm system. We established 210 annual plant communities that were subject to disturbances, fit a process-based model to their dynamics, and used counterfactual simulations to disentangle mechanisms under early-season clipping during vegetative growth and late-season clipping during reproduction. Early clipping increased diversity persistently, whereas late clipping caused a transient decline. Intrinsic demographic rates were not reduced by disturbance, and the effect of this pathway ultimately reduced diversity. By contrast, clipping weakened heterospecific competition under both treatments, thereby increasing diversity. Nearly all this increase arose from weaker effects of two dominant grasses on other species. Moreover, almost the entire diversity response to disturbance was encapsulated by these competitive effects of grasses, together with their own dynamics. Thus, diversity responses emerge from opposing demographic and competitive changes, whose balance determines whether diversity increases or declines.

## 1 Introduction

Over millennia, humans have transformed ecosystems worldwide through physical disturbance, removing plant biomass by cutting, mowing, and livestock grazing [1, 2]. Today, land-use change alters the intensity, frequency, and timing of these disturbance regimes on a global scale [3–5]. Accordingly, disturbance occupies a central place in ecological theory, particularly in explanations of species diversity [6–8]. Its negative effects are intuitive: by removing biomass, disturbance can reduce population sizes and increase the risk of local extinction. Yet disturbance can also increase diversity, particularly at intermediate levels of intensity or frequency [8], and such increases are widely observed [9, 10], especially in grasslands [11–13].

The mechanisms underlying this diversity increase have been controversial, with many competing explanations [14, 15]. Among them, two particularly influential hypotheses were introduced several decades ago and continue to shape the disturbance-diversity debate. Huston’s intrinsic demographic hypothesis [7, 15] posits that by reducing reproduction and increasing mortality, disturbance lowers intrinsic population growth rates. Slower intrinsic growth delays competitive exclusion and can therefore maintain higher diversity transiently and, when local extinction is balanced by colonization, at equilibrium [16, 17]. In contrast, Grime’s general competition hypothesis [6] proposes that disturbance weakens competitive interactions. The resulting competitive release can reduce competitive exclusion and therefore increase diversity. Distinguishing these mechanisms is important for predicting when disturbance maintains diversity and when it instead accelerates species loss.

Although many studies have documented diversity responses to disturbance, much of this work has focused on the patterns of disturbance–diversity relationships, including unimodal responses and their dependence on productivity [18–23]. Yet similar community-level patterns can arise from the demographic and competition hypotheses, as well as from other mechanisms. Such patterns therefore do not distinguish the processes through which disturbance alters diversity [24].

The first step to distinguish these mechanisms is to express the classical hypotheses, which are largely verbal, within the standard quantitative framework of multispecies community dynamics [25–27]. To our knowledge, these hypotheses have not previously been given an explicit representation within this framework. Under our formulation, the demographic hypothesis predicts that disturbance reduces intrinsic growth rates, whereas the general competition hypothesis predicts that it weakens heterospecific competitive interactions. The simple theoretical analyses (Box 1; SI Section 1) support the core logic of both hypotheses: uniform reductions in intrinsic growth can slow competitive exclusion, while uniform reductions in heterospecific competition can increase diversity. Nonetheless, the quantitative framework also exposes important complications. Species and interactions may respond unequally to disturbance, and the diversity consequences can depend on which species and interactions are affected. In particular, weakening the effects of dominant competitors on subordinate species may increase diversity, whereas changes in other interactions may have very different consequences. In grasslands, this asymmetric scenario is especially relevant because diverse forb assemblages are often suppressed by dominant grasses and released following biomass removal [28–30].

Accordingly, support for either hypotheses requires two tests: whether disturbance changes the relevant parameters in the predicted direction, and whether those changes have the predicted consequences for diversity (Fig. 1). Importantly, this framework makes both these core predictions and the more nuanced possibilities empirically testable. Disturbance effects on demographic and competitive parameters can be estimated across species and interactions, while counterfactual simulations can test whether these changes have the predicted consequences for diversity and further disentangle the contributions of particular species, interactions, and combinations of pathways.

**Figure 1:**
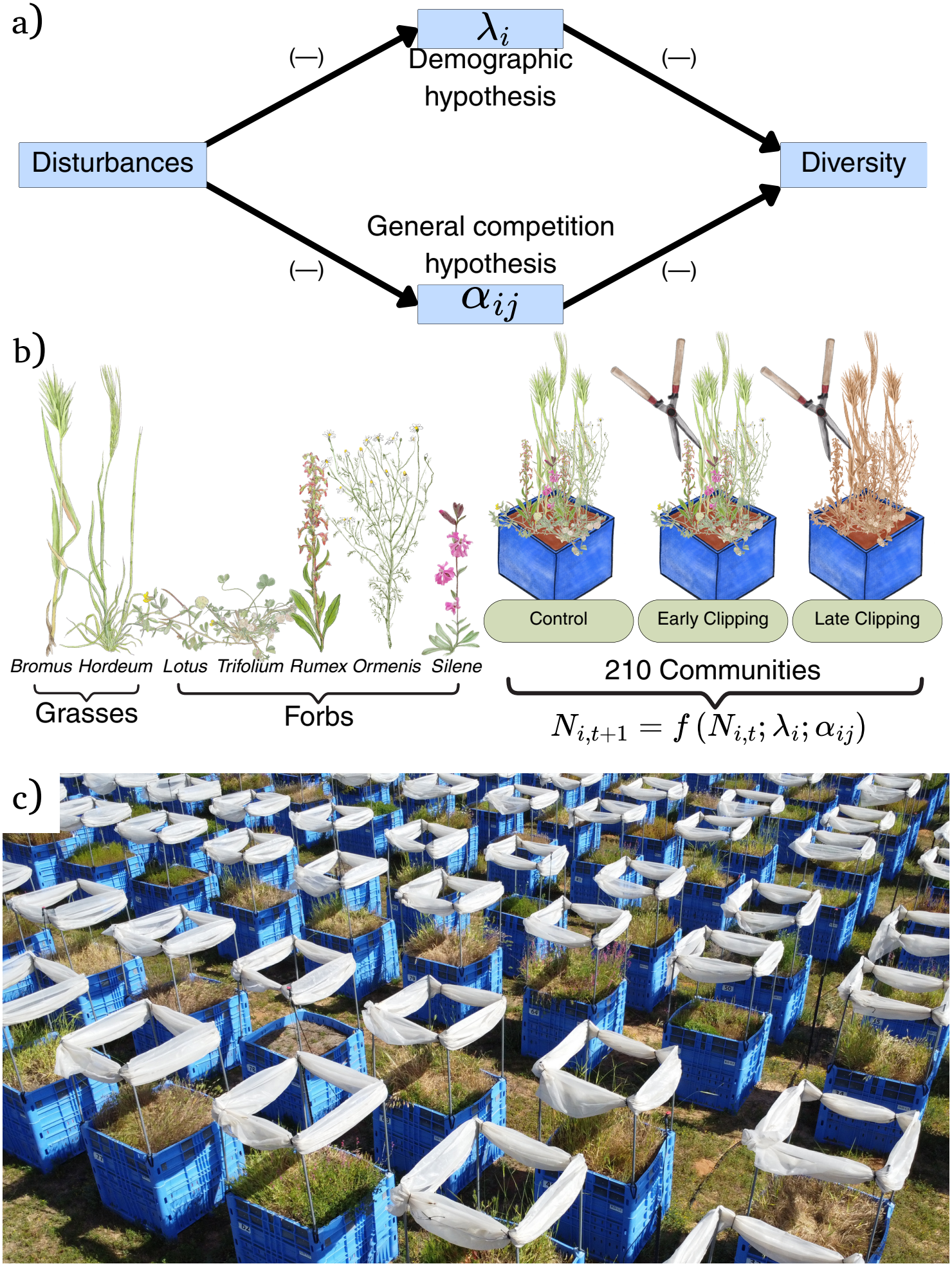
Experimental test of demographic and competitive pathways linking disturbance to species diversity. **a)** Causal diagram summarizing the two hypothesized pathways linking disturbance to species diversity. A negative sign on an arrow indicates that an increase in the first variable reduces the variable it points to. The intrinsic demographic and general competition hypotheses propose two different pathways by which disturbance can increase diversity. Under the demographic hypothesis, higher intrinsic demographic rates accelerate competitive exclusion, so disturbance can maintain diversity if it reduces these rates and thereby slows exclusion. Under the general competition hypothesis, stronger heterospecific competition promotes exclusion, so disturbance can maintain diversity if it weakens interactions among species. Thus, in both pathways, two negative effects combine into a positive net effect of disturbance on diversity. **b)** Experimental communities comprised seven annual plant species, two grasses and five forbs (i.e., non-grass herbs, including legumes; see the full scientific names in Methods 4.1). Communities were subjected to no clipping, early-season clipping, or late-season clipping and censused over four consecutive years. Community dynamics were described using an annual-plant competition model in which population size (*N*_*i*_) depends on intrinsic demographic rates (*λ*_*i*_) and competitive interactions (*α*_*ij*_) **c)** Aerial view of the experimental system.

Field-parameterized dynamic models are increasingly used to study species coexistence and diversity across environmental conditions [31–35]. However, applications to disturbance remain scarce [but see 36], and reliable inference is challenged by parameter identifiability, model-structure dependence, and uncertainty propagation [37]. Furthermore, because any tractable model omits some ecological processes, a key test of its adequacy is whether it reproduces the observed population dynamics, which in annual plants requires data spanning multiple generations [27].

Here, we leverage an extensive system of 210 experimental annual-plant communities to directly test the intrinsic demographic and general competition hypotheses (Fig. 1). The system combines replicated monocultures and multispecies communities spanning strong variation in species abundances, providing information for estimating intrinsic demographic parameters and competitive interactions. We followed species abundances over four consecutive years and experimentally imposed two disturbance treatments: early-season clipping during vegetative growth and late-season clipping during flowering and seed production. Because disturbance at different stages of the annual life cycle can alter both intrinsic demography and competitive interactions, these treatments allowed us to evaluate the two hypothesized mechanisms under biologically distinct forms of disturbance. We parameterized a dynamic community model and used counterfactual simulations to quantify the consequences of changes in each process for species diversity.

### Box 1.

**Quantitative Formulation of Classical Disturbance–Diversity Hypotheses**

The classical hypotheses were not formulated as complete multispecies dynamical models, so expressing them in demographic and competitive terms requires assumptions. We use the scaled Lotka–Volterra model as a minimal benchmark; alternative parameterizations and functional forms are considered in SI Section 1. For *S* competing species,

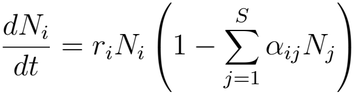

where *N*_*i*_ is the abundance of species *i, r*_*i*_ is its intrinsic growth rate (*r*_*i*_ = log *λ*_*i*_), and *α*_*ij*_ is the competitive effect of species *j* on species *i*.

**Demographic hypothesis**. If disturbance uniformly reduces intrinsic growth, 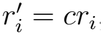, with 0 *< c <* 1, the entire right-hand side is multiplied by *c*. Competitive dynamics therefore proceed more slowly toward the same deterministic equilibrium, delaying exclusion. This directly supports the core logic of Huston’s demographic hypothesis, although the equilibrium itself is unchanged. Changes in intrinsic demography can nevertheless affect steady-state diversity through extinction–colonization dynamics or, in other population-dynamic models, through changes in equilibrium abundances (SI Section 1).

**General competition hypothesis**. The competition hypothesis concerns whether weaker competitors can persist. In the simplest two-species case, this is assessed by asking whether each species can increase when rare in a population dominated by the other. If a rare species has positive population growth, it can invade the resident population rather than being driven to extinction.

Consider species *i* when rare and species *j* at its single-species equilibrium, 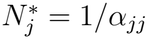. Species *i* increases when rare if

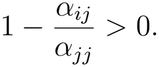

Thus, the competitive effect of resident species *j* on rare species *i* must be weaker than the self-limitation experienced by *j*. An analogous condition applies to species *j* invading species *i*. Reducing heterospecific competition while leaving conspecific competition unchanged therefore makes persistence of both species more likely and can promote coexistence.

This also shows why unequal responses matter. Weakening the effects of a dominant competitor on a subordinate species can allow the subordinate species to persist and increase diversity, whereas weakening other interactions may have little benefit or even favor the stronger competitor. The outcome therefore depends on which interactions are affected.

Overall, the classical logic holds under simple conditions, but predictions become less straightforward when species and interactions respond unequally or both pathways operate simultaneously (SI Section 1). Our empirical analysis therefore estimates species- and interaction-specific disturbance responses and uses counterfactual simulations to determine their individual and combined consequences for diversity.

## 2 Results

### 2.1 Disturbance regime alters community dynamics and diversity across timescales

The experimental communities reproduced a widely observed grassland response to disturbance. In the absence of disturbances, experimental communities became strongly dominated by the two grasses, *Bromus* and *Hordeum*, while most forbs were lost from most communities. Early-season clipping altered this trajectory and allowed several forbs to persist, consistent with the common release of subordinate forbs following biomass removal, whereas late-season clipping intensified *Bromus* dominance and suppressed *Hordeum* (Fig. 2).

**Figure 2:**
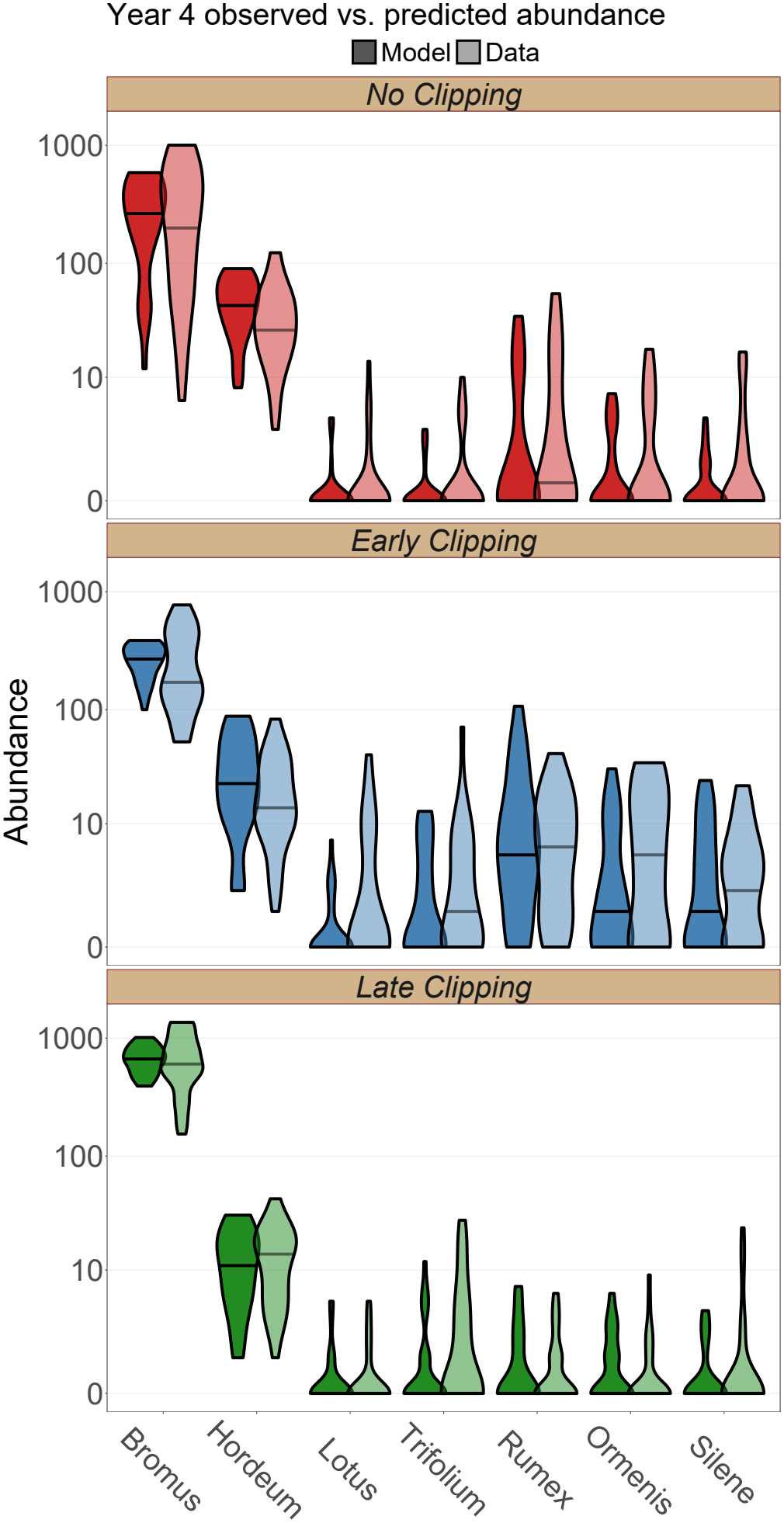
Disturbance changes species persistence, and the model reproduces these community responses. Light-colored violins show the central 90% distribution of the observed data, and dark-colored violins show the central 90% distribution of draw-level medians from the posterior predictive distribution (PPD). Horizontal lines indicate medians; where a line is not visible, the median is 0. Without clipping, the two grasses dominate and most forb species are absent from most communities. Early clipping increases forb persistence across communities, whereas late clipping decreases *Hordeum* and increases *Bromus*.

Notably, the field-parameterized model closely reproduced the observed responses at both levels of organization: species-level abundance patterns (Fig. 2) and community-level differences in diversity (Fig. 3a), with additional diagnostics and model comparisons supporting its fit (SI 2.2).

**Figure 3:**
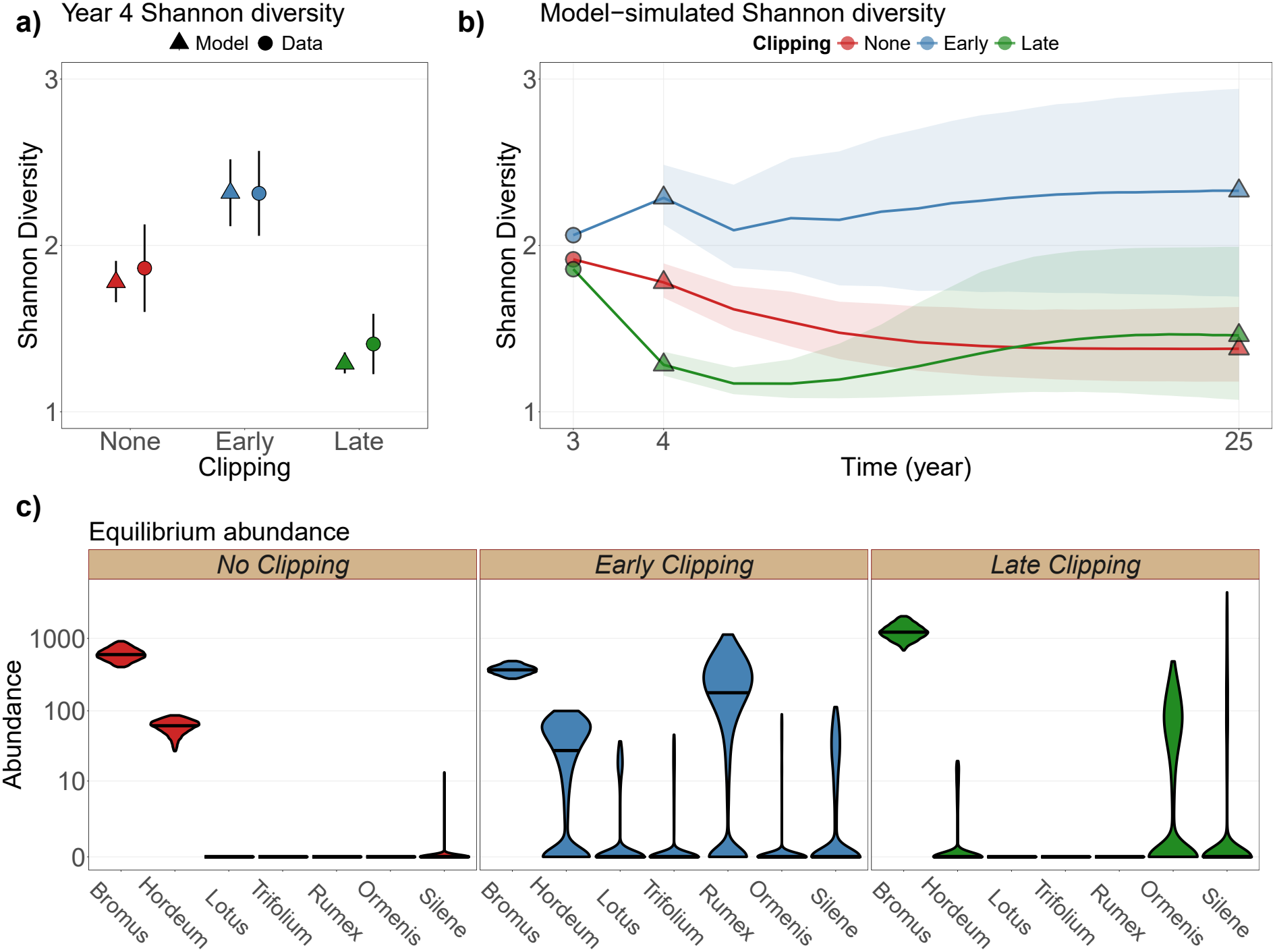
Early disturbance maintains higher diversity across timescales, while late disturbance causes a transient diversity decline. Diversity is expressed as the effective number of species, exp(*H*^′^), where *H*^′^ is Shannon entropy. **a)** Observed year-4 diversity and corresponding in-sample posterior expectations. Points represent means, and error bars represent 90% confidence intervals for the data and 90% credible intervals for the model. The model reproduces the contrasting year-4 diversity responses among clipping regimes. **b)**Deterministic model projections initialized with observed population sizes from years 2 (*t* − 1) and 3 (*t*; circles), shown on a logarithmic time scale. The diversity increase under early clipping persists to equilibrium, reached approximately by year 25, whereas diversity under late clipping partially recovers and converges with the control. Paired posterior contrasts strongly support higher equilibrium diversity under early clipping than under either the control or late clipping (Fig. S4a). Triangles represent posterior medians at selected time steps, and shaded regions represent 90% credible intervals. **c)** Equilibrium species abundances under each clipping regime. Control communities are dominated by the two grasses. Early clipping allows *Rumex* to persist and become the second-most abundant species, whereas under late clipping *Bromus* dominates, *Hordeum* is nearly extinct, and *Ormenis* becomes the second-most abundant species. Violins show the central 90% of the distribution of draw-level medians; horizontal lines indicate medians; where a line is not visible, the median is 0.

These contrasting community trajectories among treatments translated into different diversity responses across timescales (Fig. 3). Early clipping increased diversity during the experiment, and this increase persisted to deterministic equilibrium (equilibrium is reached by year 25, see Fig. S2). Late clipping instead reduced diversity during the experiment, but diversity subsequently recovered and converged with the control at equilibrium (Fig. 3a,b). The temporal diversity responses were qualitatively similar when using Simpson diversity (Fig. S3). Despite this convergence in diversity, late clipping produced a distinct equilibrium composition, as did early clipping (Fig. 3c).

### 2.2 Disturbance has contrasting effects on competition and intrinsic demography

In contrast with the prediction of the demographic hypothesis, disturbance did not generally reduce intrinsic growth rates, *λ*_*i*_ (Fig. 4a,b). *Bromus*, which had the highest baseline *λ*_*i*_ (Fig. 4a) and dominated the experimental communities, showed an increase rather than a decrease under disturbance (Fig. 4b). Only *Hordeum* showed a modest decline, while responses among the remaining species were weak or positive (Fig. 4b). An additional lagged demographic term, *s*_*i*_, likewise showed no consistent reduction under disturbance (Fig. S5b). Thus, disturbance did not produce the systematic decline in intrinsic demography predicted by the demographic hypothesis.

**Figure 4:**
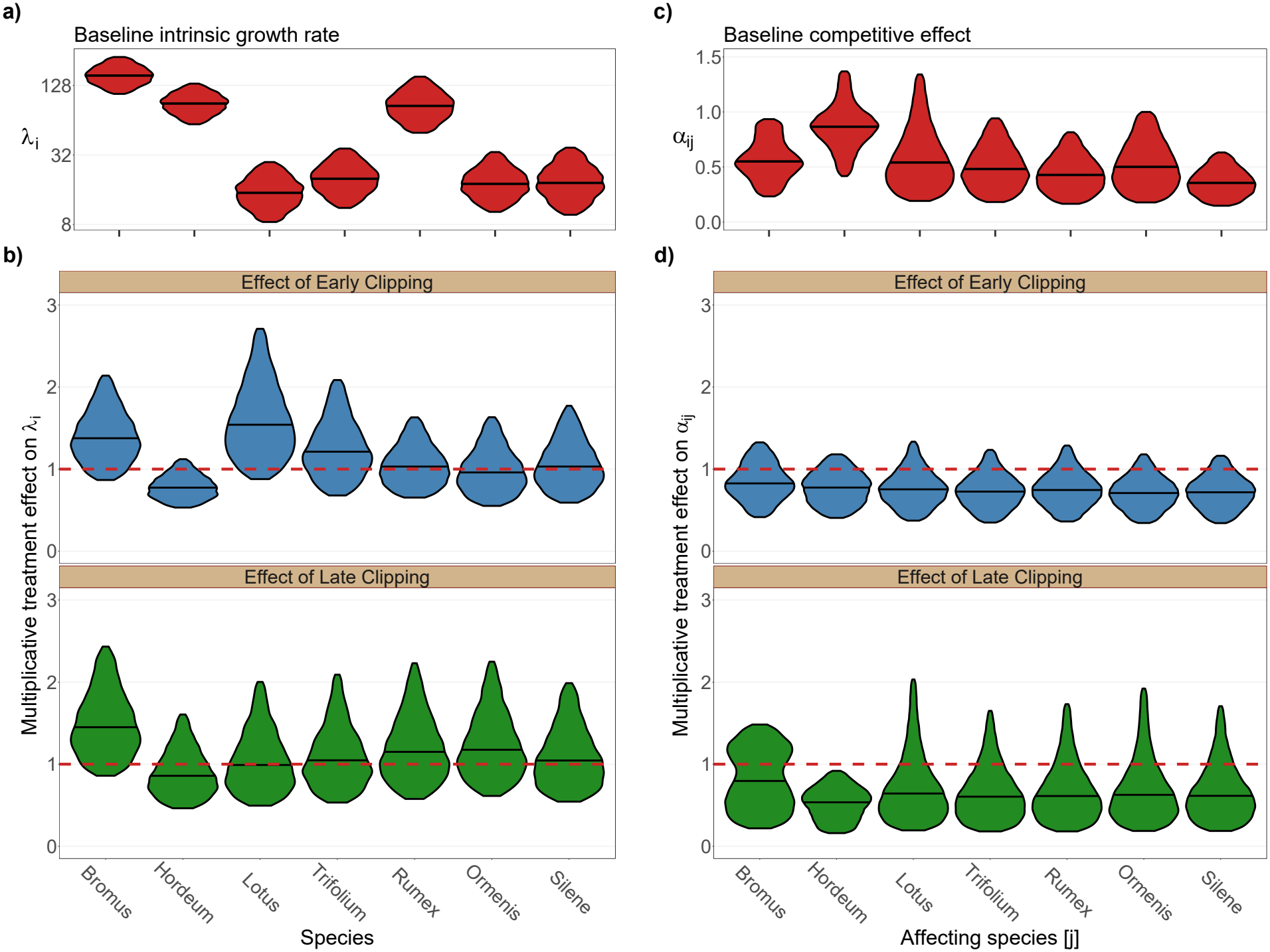
Disturbance weakens heterospecific competition but does not reduce intrinsic growth rates. Violins show the central 90% of posterior distributions, with horizontal lines indicating medians. **a)** Baseline intrinsic growth rate, *λ*_*i*_ (the first term on the RHS in Eq.S10), for each of the seven experimental species in control plots, shown on a logarithmic scale. *Bromus* has the highest baseline intrinsic growth rate. **b)** Multiplicative effect of each clipping regime on *λ*_*i*_ (the second term on the RHS in Eq.S10). The dashed red line indicates no treatment effect; values above 1 indicate an increase under clipping and vice versa. Clipping does not generally reduce intrinsic growth rates and instead increases them for several species, particularly the dominant *Bromus*. **c)** Baseline heterospecific competitive effects, *α*_*ij*_ (the first term on the RHS in Eq.S13), in control plots. Species names denote the competitor species *j*, and competitive effects are pooled across all focal species *i* ≠*j*. Thus, for example, the value shown for *Bromus* summarizes the effects exerted by *Bromus* on the other species. *Hordeum* exerts the strongest baseline competitive effects. **d)** Multiplicative effect of each clipping regime on heterospecific competition (the second term on the RHS in Eq.S13), summarized by competitor species *j* as in **c)**. The dashed red line indicates no treatment effect.

In accordance with the general competition hypothesis, disturbance weakened heterospecific competitive effects under both clipping regimes across all species (Fig. 4d). Baseline competitive effects differed among species, with *Hordeum* exerting the strongest effects (Fig. 4c), but clipping broadly reduced these effects regardless of species identity (Fig. 4d). The same general pattern emerged for competitive sensitivity, with most species becoming less sensitive to heterospecific competition under disturbance (Fig. S6b). In contrast, conspecific competition showed no comparable systematic response (Fig. S7b).

### 2.3 Opposing demographic and competitive pathways determine diversity responses

Counterfactual simulations tested the diversity consequences of the two general mechanisms by retaining disturbance effects only on the parameter subset associated with each hypothesis. To implement these simulations we used the model-estimated parameters and simulated the communities to equilibrium, but turned on the effect of disturbance only on the parameter subset tested (Fig. 5). When disturbance acted only through intrinsic demographic parameters, *λ*_*i*_ and *s*_*i*_, equilibrium diversity decreased under both clipping regimes (Fig. 5a, in this figure, compare the green and blue triangles, representing the counterfactual disturbance treatment, to the red control). In contrast, when disturbance acted only through heterospecific competition, diversity increased under both regimes (Fig. 5b). Thus, disturbance-induced changes in intrinsic demography and competition had opposing consequences for diversity.

**Figure 5:**
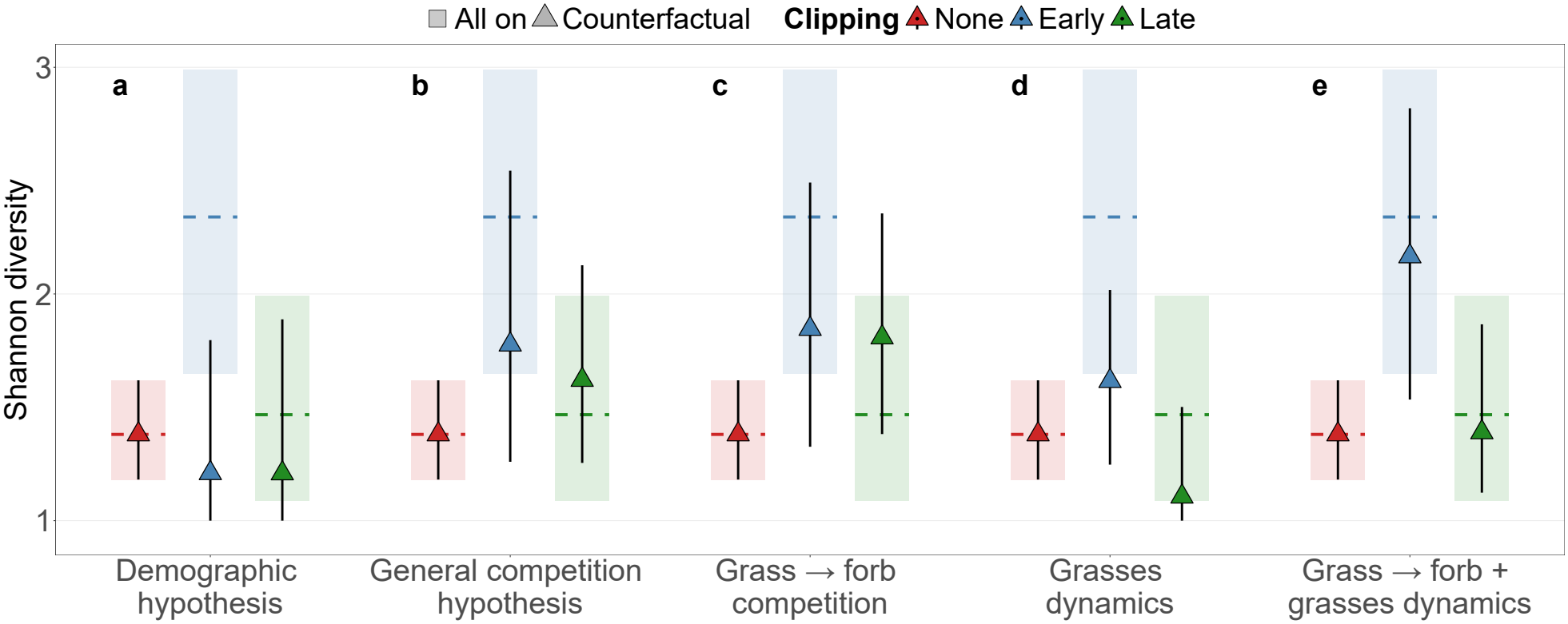
Counterfactual simulations separate demographic, competitive, and dominant-grass contributions to equilibrium diversity. Simulations were run to equilibrium and initialized with observed abundances from years 2 (*t* − 1) and 3 (*t*). Rectangles and dashed lines show posterior medians and 90% credible intervals, respectively, for the full model, in which disturbance affects all parameters; these are identical across panels **a–e**. Triangles and error bars show posterior medians and 90% credible intervals for counterfactual simulations in which disturbance effects are retained only for the parameter subset specified in each panel. Diversity is expressed as the effective number of species, exp(*H*^′^), where *H*^′^ is Shannon entropy. **a)** Disturbance effects retained only for the intrinsic demographic rates *λ*_*i*_ and *s*_*i*_. **b)** Disturbance effects retained only for all 42 heterospecific competitive interactions. **c)**Disturbance effects retained only for the 10 grass → forb interactions. The resulting diversity increase is similar to that obtained when all heterospecific interactions are modified in **b. d)** Disturbance effects retained only for parameters governing the two grasses’ intrinsic demographic rates, conspecific regulation, and competition with each other. **e)** Disturbance effects retained for the dominant-grass parameters in **d**, together with (as in in **c**) grass → forb competition. This combined subset reproduces nearly the full diversity response of the complete model using 18 of the 63 dynamic parameters.

Next, we tested the hypothesis that diversity increases through competitive release of forbs from dominant grasses. Restricting disturbance effects to the 10 grass → forb interactions produced nearly the same diversity increase as modifying all 42 heterospecific interactions (Fig. 5b,c), whereas disturbance effects restricted to competition exerted by forbs had a negligible effect on diversity (Fig. S8). Thus, although disturbance broadly weakened heterospecific competition, its positive effect on diversity stemmed primarily from reducing the competitive pressure exerted by grasses on forbs..

Because community dynamics were strongly dominated by *Bromus* and *Hordeum*, we also tested whether disturbance effects involving these two species could explain the contrasting responses to early and late clipping. Restricting disturbance effects to parameters governing the grasses’ intrinsic demographic parameters, conspecific regulation, and competition with each other reproduced the direction of the contrast between regimes, with higher diversity under early than late clipping (Fig. 5d; Fig. S9). However, diversity under both clipping regimes remained substantially lower than in the full model (triangles and error bars, representing counterfactual simulations, have lower diversity than the rectangles, representing the full model in Fig. 5d), because the diversity-promoting reduction in grass → forb competition was absent.

Finally, combining these grass-dynamics effects with grass → forb competition reproduced nearly the full diversity response of the complete model while involving only 18 of its 63 dynamic parameters (Fig. 5e). Under early clipping, the diversity benefit of weakened competition outweighed the opposing grass-dynamics effects, whereas under late clipping it did not. The counterfactual results were qualitatively unchanged for Simpson diversity (Fig. S10).

## 3 Discussion

Early- and late-season clipping produced contrasting diversity responses, yet both altered the processes proposed by the two hypotheses. The demographic hypothesis received only partial support: higher intrinsic demographic rates reduced diversity as predicted, but clipping generally did not reduce these rates, so changes in intrinsic demography tended to decrease rather than increase diversity. In contrast, heterospecific competition weakened under both clipping treatments, supporting the general competition hypothesis, and this weakening increased diversity, although the resulting diversity increase resulted primarily from the effects of the dominant grasses on the rest of the community. Changes involving these dominant species also captured much of the contrasting diversity response to early and late clipping.

The unexpected increase in demographic rates may partly reflect the distinction between reproductive output and whole-life-cycle population growth. Many plant competition studies estimate intrinsic growth from one-generation experiments that manipulate neighbor density and measure reproductive output, while treating other components of the annual life cycle, such as seed survival and germination, as fixed [36, 38, 39]. In contrast, our approach of repeated population censuses allows *λ*_*i*_ to be estimated from realized annual population transitions [40], thereby integrating reproduction, seed survival, germination, and establishment into a single measure of potential population growth. As with any quantitative expression of a largely verbal hypothesis, alternative formalizations are possible, but our interpretation is closely aligned with Huston’s explicit emphasis on disturbances that “reduce growth rates and the rate of competitive exclusion” [15, p.2391]. Reduced biomass or fecundity after disturbance therefore need not imply lower *λ*_*i*_ if reproductive losses are offset elsewhere in the life cycle, for example through compensatory growth or improved recruitment. Indeed, responses were species-specific: disturbance modestly reduced *λ*_*i*_ in the tall, later-phenology *Hordeum*, had little effect on several species, but increased it in *Bromus*, particularly under late clipping.

Disturbance consistently weakened heterospecific competition across species, providing stronger support for the general competition hypothesis than expected. The reduction under both clipping regimes is consistent with different biological mechanisms. Early clipping, imposed during active vegetative growth, reduces canopy size and can thereby alleviate asymmetric competition for light [11, 13, 29]. Late clipping, imposed near the end of the growing season, may additionally reduce the standing biomass and litter carried into the following generation, thereby alleviating inhibition of seedling recruitment [41, 42]. Crucially, nearly the entire competition-mediated increase in diversity was reproduced by weakening the effects of the two dominant grasses on forbs, consistent with the long-standing hypothesis that diversity increases when subordinate species are released from dominant competitors [30, 43–46]. The two grasses may influence diversity through different routes: *Hordeum* exerted the strongest per-capita competitive effects, whereas *Bromus* reached the highest abundance, so even more moderate per-capita effects could translate into a large community-level impact.

The role of the dominant grasses extended beyond their competitive effects on forbs. Because forbs comprised the majority of species in our communities, as they commonly do in grasslands, one might expect changes in their demography and interactions with one another to dominate the diversity response. Instead, these processes contributed little to the community-level pattern. Disturbance effects on the intrinsic demography, conspecific regulation, and reciprocal competition of the two dominant grasses reproduced the direction of the contrast between early and late clipping, and adding their competitive effects on forbs nearly reproduced the full diversity response using only 18 of the 63 dynamic parameters. The contrast between disturbance regimes also depended on species-specific changes in conspecific regulation, particularly for *Bromus* (Fig. S9), indicating that the two original hypotheses capture major pathways of disturbance effects but not every component of the community response. Notably, this disproportionate role of dominant species is consistent with recent global evidence that changes in dominant species strongly predict changes in grassland species richness [47].

Our approach is in line with recent empirical applications of coexistence theory, which typically combine two complementary elements: decomposition of population growth into theoretically defined quantities, such as niche and fitness differences or fluctuation-dependent and independent mechanisms, and counterfactual simulations that quantify the contribution of particular processes [48, 49]. Here, our hypotheses concerned intrinsic population growth and competitive effects, rather than niche and fitness differences or other standard coexistence quantities, so we did not apply the first type of decomposition. Instead, we used the second element, combining a fitted population model with counterfactual simulations to isolate the consequences of the specific processes implicated by the hypotheses. Fitting explicit population dynamics also allowed the same inferred processes to be examined across the transient timescale of the experiment and their longer-term deterministic tendencies. Our experiment was designed around mechanisms operating under fixed disturbance regimes, but the same general strategy could be extended to disturbance mechanisms that depend explicitly on spatial or temporal variability, such as regeneration niches, competition-colonization trade-offs, or temporally varying disturbance, provided that the relevant demographic, dispersal, or temporal processes are represented in the experiment and population model.

This mechanistic perspective is particularly relevant, as human activities are changing not only the frequency and intensity of disturbance, but also its timing within the life cycle of organisms. Our results show that such changes in disturbance regime can produce contrasting diversity outcomes even when the physical disturbance is similar. Predicting biodiversity responses in a changing world therefore requires a mechanistic understanding of how disturbance regimes affect the demographic and competitive processes underlying community dynamics.

## Author Contributions

The research questions and conceptual framing of this study were developed by Y.N., M.K. and N.D.; the experimental system was designed by N.D., O.G. and T.S.; experimental setup and sampling were performed by Y.N., O.G. and T.S. Analyses were designed and performed by M.K. and Y.N., with input from N.D.; Y.N., N.D. and M.K. wrote the first draft of the manuscript, and all authors contributed to subsequent revisions. N.D. and M.K. jointly supervised the work.

## Acknowledgements

We thank Sara Barcza, Gal Ben Shlomo, Noa Levi, Avishag Neeman, Yarden Schindler and Shlomo Shik for their assistance in fieldwork, Sara Barcza for the illustrations in Figure 1, and Ronen Kadmon for valuable comments and suggestions on the manuscript.

## Funding

This study was funded by the Israel Science Foundation (grants no. 2403/22, 672/22 and 2426/24) to N.D and M.K. Y.N. and T.S. were supported by the Women’s League for Israel. Y.N. and O.G. were supported by the School of Environmental Studies of the Hebrew University, Hebrew University Center for Sustainability. Y.N., T.S. and O.G. were supported by the Jewish National Fund.

## Data and Code Availability

All the raw data files, along with all custom scripts needed to fully reproduce the results presented in this manuscript, can be found in https://doi.org/10.5281/zenodo.24049074.

## 4 Methods

### 4.1 Experimental system

The experiment was designed as a simplified representation of grassland communities in which species compete over successive generations, and physical disturbance can alter those dynamics. We therefore assembled communities containing grasses together with a diverse set of forbs, including legumes, capturing a common structural feature of grasslands: grasses often dominate community abundance, whereas forbs account for much of plant species diversity and commonly benefit from biomass-removing disturbance [28, 30]. We used plant communities of the red-sandy soils of Israel’s coastal region, hereafter Hamra communities, as a model system for these dynamics. These communities have a long history of anthropogenic disturbance, including grazing and mowing, and are often dominated by annual plants [50]. The combination of discrete annual generations and high local population densities, often reaching thousands of individuals per square meter, provides a relatively tractable system for estimating whole-community population dynamics and species interactions across successive generations.

The mesocosm experiment was established during the summer of 2021 at the experimental farm of the Hebrew University of Jerusalem in Rehovot, Israel (31.906268, 34.800049), with elevation ≈ 44 m ASL. The region has a Mediterranean climate, with cool, rainy winters and hot, dry summers, and mean annual rainfall of approximately 540 mm year^−1^ (Table S1).

The experimental species pool comprised seven common local annual species: two grasses (*Poaceae*), *Bromus scoparius* and *Hordeum spontaneum*, two legumes (*Fabaceae*), *Lotus halophilus* and *Trifolium tomentosum*, and three other forbs, *Rumex bucephalophorus, Ormenis mixta* and *Silene palaestina*. This restricted species pool retained the major functional groups relevant to the grassland structure we aimed to represent, while allowing the abundance and interactions of every species to be followed through time.

Communities were established in cuboid plastic containers (1.035 m × 1.035 m × 1 m) filled with local Hamra soil, providing sufficient soil depth for root development and water storage. Each container was fitted with 1 m high poles supporting a rolling net. Nets were kept open during the growing season to allow airflow, maintenance, and sampling, and closed toward the end of the season to minimize seed dispersal and migration among communities.

To generate strong variation in conspecific and heterospecific abundances, species were established in seven monocultures and seven multispecies initial compositions. In each mixture, one species was targeted to comprise 60% of seedlings and each of the other six species 6.7%. Seed numbers were adjusted using preliminary germination trials to approximate these target frequencies while maintaining a similar total seedling density among communities; subsequent analyses used the observed population densities rather than the target frequencies. Monocultures were maintained by removing all other species, and nonexperimental species were continuously removed from both monocultures and mixtures. This maintained a defined species pool while allowing the experimental species to undergo natural changes in relative abundance through time.

Communities were assigned to one of three clipping regimes using complete randomization, without spatial blocks. Clipping treatments were applied annually from the second growing season onward, after communities had established. At each clipping event, all vegetation was cut to 7 cm and the removed biomass was discarded, mimicking biomass removal by cattle grazing typical of Mediterranean grasslands [46]. Early-season clipping was applied during vegetative growth once the tallest plant in a random sample of 10 communities reached 40 cm, usually in late December. Using a height threshold rather than a fixed date ensured that clipping occurred at a comparable stage of canopy development despite interannual variation in germination and growth. Vegetation, particularly the grasses, regrew rapidly during winter, and communities were clipped again after reaching the threshold height approximately 3–4 weeks later. Late-season clipping was applied once in early April, during flowering and seed production, when some species had already set seed and senesced while others were beginning to flower. A second late-season clipping was not biologically meaningful because vegetation subsequently senesces during the dry Mediterranean summer and does not regrow before the next growing season. Control communities remained unclipped.

The experiment therefore comprised 14 initial-composition treatments, seven monocultures and seven mixtures with a different initially dominant species, each exposed to the three clipping regimes and replicated five times, for a total of 210 communities:

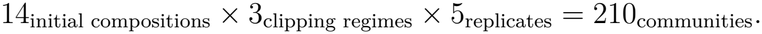

Species abundances were censused annually during peak biomass in March–April, before the late-season clipping treatment was applied. Within each container, all individuals were counted in a permanent central 40 cm × 40 cm quadrat. Sampling the center of the container reduced edge effects while allowing the same communities to be followed nondestructively through time. Because seeds produced outside the central quadrat could subsequently recruit within it, the population model included a lagged demographic term that can account for both seed-bank contributions and within-container edge-to-center recruitment (see 4.2.1). Further details of experimental setup and maintenance are provided in SI 2.1.

### 4.2 Field-parameterized model

For each plot-year combination, species counts were summed across the four squares to obtain plot-level population sizes. The data were then arranged as a data frame in which each row represented a distinct plot × year combination, with each species’ abundance given at times (*t*−1), (*t*), and (*t*+1). Details of incomplete counts, abundance corrections, and plot exclusions are provided in SI 2.2.1. The model was implemented using the `Stan` language and fitted in R v4.6.0 [51], using the `rstan` package [52]. Absolute goodness-of-fit was tested using the `DHARMa` package [53] and by simulating the model to project an observed time step using the previous ones (Fig. 2) [27], while relative goodness-of-fit compared to other functional forms was considered using the loo package [54]. Full sampling settings, convergence criteria, and diagnostic procedures are provided in SI 2.2. All data manipulations, as well as visualizations, were performed with `tidyverse` packages [55].

#### 4.2.1 Functional form

For our model, we chose to use the functional form proposed by Law & Watkinson [56]:

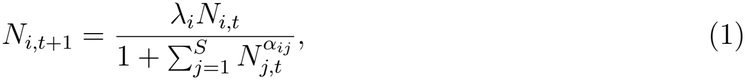

where *N*_*i,t*_ is population density for species *i* at time *t, λ*_*i*_ is that species’ maximal growth rate in the absence of competitors, *α*_*ij*_ is the effect of species *j* on species *i*, and *S* is the total number of species.

A few changes were made to the original form: a) the effects of disturbance regimes and temporal variability on *λ*_*i*_ and *α*_*ij*_ were incorporated into the model (see 4.2.2); b) to account for the discreteness of individuals and to ensure *λ*_*i*_ represents growth rate when one individual is present, the conspecific population-density term *N*_*i*_ was replaced with the number of neighbors, *N*_*i*_ − 1; and c) to incorporate our prior knowledge of seed bank dynamics in grasslands [57, 58], as well as the possibility of edge → center within-plot immigration, a lagged response parameter *s*_*i*_ multiplying the population size of species *i* from year *t* − 1 was incorporated; this parameter is also affected by disturbance and temporal variability. Thus, our final model form is:

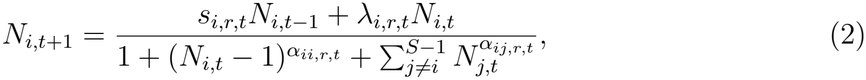

where all parameters are identical to Eq. (1), with the addition of *s*_*i*_ as the seed bank contribution maximal growth rate, i.e., the number of individuals at *t* + 1 when at *t* − 1 there was only one individual and at *t* population size was zero. The subscripts *r* and *t* indicate the disturbance regime and experimental year, respectively (see SI 2.2.3 for further details on the hierarchical structure of the parameters and their dependence on year and treatment).

This modified form was chosen using the `loo_compare()` function from the loo package [54, 59] against other classical functional forms [60, 61] and modifications. The observed population sizes were assumed to follow a negative binomial distribution around their expected sizes (given by Eq. 2), using an altered noise parameterization (see SI 2.2.2 for details). Full model-comparison procedures and results are provided in the SI, section 2.2.1, and Tables S2 and S3. Full information on priors is provided in SI 2.2.4.

#### 4.2.2 Clipping & year effects

We tested two major hypotheses that can be viewed as predictions on disturbance’s effect on parameters of community dynamics; hence we incorporated a multiplicative treatment parameter for each of the parameters in Eq. 2 to quantify this effect. The complete parameterization is provided in SI 2.2.3.

Mediterranean systems are known for high temporal variability in climatic conditions [62], and our system showed high variability in both precipitation and temperature (Table S1). To account for this, we hierarchically modeled the effects of specific years on each parameter in every species. We assumed the geometric mean 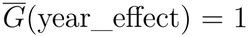, so that year effects represent temporal deviations around the expected dynamics. The biological logic for the year effects is presented in SI 2.1, and their hierarchical parameterization is provided in SI 2.2.

### 4.3 Simulations

To understand the ultimate state of the system under different clipping regimes, we wanted to quantify diversity at the deterministic equilibrium. We simulated community dynamics by iteratively applying Eq. (2) with the posterior parameter values estimated for each species. For each posterior draw and treatment, expected abundances at time *t* + 1 were calculated from abundances at time *t* and *t* − 1, and then used as the starting point *t* for the next time step. Simulations were run for 1 × 10^3^ discrete generations and initialized with observed abundances from years 2 (*t* − 1) and 3 (*t*). In every scenario we simulated the 104 experimental mixture communities included in the analysis (SI 2.1), each with that community’s observed species abundances at years 2 (*t* − 1) and 3 (*t*) as initial conditions. Every community was simulated 5.4 × 10^3^ times, each with a different posterior draw, using only the time-invariant parameters (i.e. excluding temporal variation). . Diversity was calculated as Shannon diversity, Simpson diversity, and species richness [63], and extracted as the median diversity for a given posterior draw in each treatment across all 104 simulated communities, providing us with three posterior distributions for each diversity index.

In order to isolate specific hypothesis-driven mechanisms, counterfactual simulations where disturbance acts only on specific parameters of community dynamics were necessary. The simulations were thus designed with a “switchboard”, allowing us to manually “switch on” the effect of disturbance on each intrinsic parameter, i.e., *s*_*i*_, *λ*_*i*_, and *α*_*ii*_. This switch can be activated for any given set of one or more parameters and species. For *α*_*ij*_, the switch allows specifying focal species *i* and competitor *j* | (*i* ≠*j*) for any specific *α*_*ij*_s. Diversity was calculated as Shannon effective number of species, and Simpson effective number of species, with abundances below 1 treated as zero. Details of simulation failures can be found in Table S4, and details on the differentiation of equilibrium diversity patterns are provided in SI 2.3.1.

The dynamic Bayesian regression model we fit is composed of a deterministic and stochastic component. While in the real world stochasticity may play an important role in shaping community dynamics, the tested hypotheses directly address the deterministic component (i.e., *λ*_*i*_&*α*_*ij*_) and thus we used it to predict equilibrium diversity patterns. Similarly, We acknowledge that non-linear averaging effects may alter the deterministic equilibrium, but since they are not part of the hypotheses we test, we did not incorporate them into the simulations of the equilibrium. Still, when making short-term predictions using both stochastic and deterministic components, we reliably replicated species-level abundance data across treatments (Fig. 2), suggesting that the model faithfully captures how the system operates [27].

## SI

### 1 Expressing verbal hypotheses as a quantitative framework

Our work addresses hypotheses that were either completely verbal [1] or mostly verbal [2] and translates them into quantitative predictions, a task that, like any translation work, requires assumptions and may allow for different interpretations. We represent species-specific demographic parameters through vectors, such as the finite rates of increase ***λ*** = (*λ*_1_, …, *λ*_*S*_)^⊤^, and competitive effects through an interaction matrix ***A*** = (*α*_*ij*_), where *α*_*ij*_ represents the competitive effect of species *j* on species *i*. Disturbance-induced changes in these parameters can then be related to changes in community diversity. We begin with Lotka– Volterra (LV) models because their mathematical simplicity facilitates the derivation of theoretical predictions and the identification of underlying assumptions, and because Huston’s original formulation relied on LV. However, these predictions do not necessarily extend unchanged to other formulations of population dynamics, including Beverton–Holt, Ricker, and Law–Watkinson models [3–6]. We therefore also derive invasion conditions for a simplified Law–Watkinson model and examine the diversity consequences of reduced competition through simulations using the seven-species Law–Watkinson model fitted to our experimental data. This same model underlies our empirical analysis of disturbance-induced changes in demographic and competitive parameters, connecting the theoretical predictions to their experimental evaluation.

#### The demographic hypothesis

Huston [2] proposed a verbal hypothesis backed with a simple proof in a two-species Lotka–Volterra simulation [3], which we solve analytically below to show it applies to any *S* number of species. The two-species ODE system is:

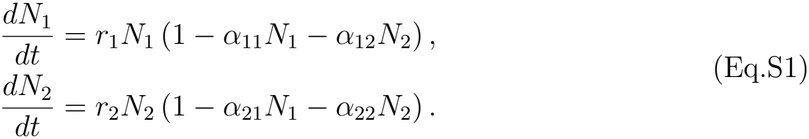

Where *N*_*i*_ is the abundance of species *i, r*_*i*_ its intrinsic growth rate (*r*_*i*_ = log *λ*_*i*_ for a unit time interval), *α*_*ii*_ represents conspecific competition, and *α*_*ij*_ (*i*≠*j*) represents the competitive effect of species *j* on species *i*. Carrying capacities are implicit in the conspecific competition coefficients, with *K*_*i*_ = 1*/α*_*ii*_.

Huston showed that uniformly reducing the intrinsic growth rates *r*_*i*_ (say, by disturbance) slows down species’ approach to equilibrium, and thus delays competitive exclusion. Showing that mathematically, let’s assume that all population growth rates are reduced by 0 *< c <* 1, such that *r*_1_ Ô→ *cr*_1_, *r*_2_ Ô→ *cr*_2_. Thus, the entire right-hand side of Eq.S1 is multiplied by *c*. Consequently,

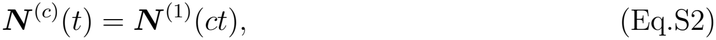

where the superscripts indicate the modified and original growth rates, respectively. This identity follows because the rate of change of ***N*** ^(1)^(*ct*) is *c* times the original rate of change. Thus, Huston’s core logic is correct, as it applies to any number *S* of species in this LV formulation. Although this slowing leaves equilibrium diversity unchanged in the LV formulation considered here, demographic changes can also affect steady-state diversity when local extinction and colonization are incorporated [7, 8].

However, two main complications arise when extending this prediction: model formulation and differential species responses. The first arises even within Lotka–Volterra, which can be expressed either as in Eq.S1 or in its unscaled form:

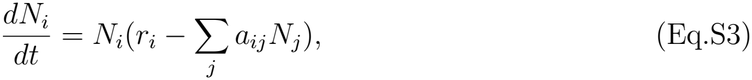

where the competition coefficients in the two formulations are related such that *a*_*ij*_ from Eq.S3 equals *r*_*i*_*α*_*ij*_ in Eq.S1. Although mathematically equivalent, reducing growth rates while keeping the respective competition coefficients fixed has different consequences. In the former, equilibrium abundances remain unchanged, whereas in the latter, reducing *r*_*i*_ also reduces carrying capacity, since *K*_*i*_ = *r*_*i*_*/a*_*ii*_ [9]. To obtain the same demographic slowing in the latter formulation, all *a*_*ij*_, including conspecific and heterospecific coefficients, must be reduced proportionally with *r*_*i*_. Similar complications arise in other density-dependent models [4–6], where reducing growth rates may also change equilibrium abundances and the competitive pressure experienced by other species.

A second complication is that demographic responses may differ among species. For two species whose growth rates are reduced by different factors, 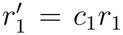 and 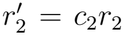, the change in their relative abundances is given by

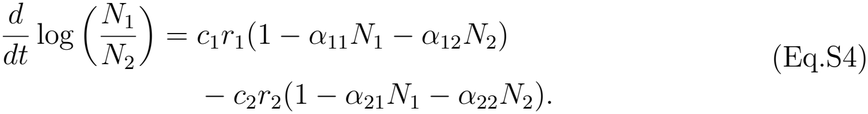

When *c*_1_ = *c*_2_, relative-abundance dynamics are uniformly slowed; when *c*_1_≠*c*_2_, the direction of relative change may also differ. Thus, equilibrium abundances remain unchanged, but the consequences for transient diversity depend on which species’ growth rates are affected and by how much.

To summarize, the “slowing down” suggested by the demographic hypothesis applies wherever disturbance reduces all rates uniformly and these rates do not affect equilibrium densities; where these are not the case, further examination of the diversity consequences of disturbance is required.

#### The general competition hypothesis

The original paper by Grime [1] presents a verbal hypothesis which relies on his observation that species richness is high in communities with a low mean ‘competitive index’ (his Figure 1). Grime relates lower species richness in communities with high competitive index to competitive exclusion, and argues that under physical disturbance, the dominant competitors “…decline in vigour and species of lower competitive ability are able to survive.” (p. 345). Based on this argument, we interpret Grime’s hypothesis as a reduction in heterospecific competitive effects that allows weaker competitors to persist. To examine the general consequences of such a reduction, we initially assume a common disturbance effect *c* across all heterospecific competition parameters, such that 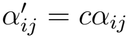 for *i* ≠*j*, where *α*_*ij*_ describes the competitive effect exerted by species *j* on species *i*, and 0 *< c <* 1.

The same two-species scaled Lotka–Volterra model (Eq.S1) illustrates this logic through the conditions for species invasion. When species *j* is at its monoculture equilibrium, 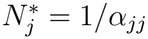, species *i* can increase when rare if [10]

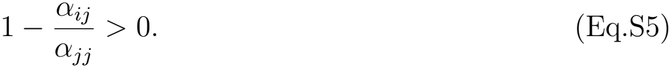

Reducing heterospecific competition while leaving conspecific competition unchanged makes this condition easier to satisfy. In contrast, reducing the entire interaction matrix by a common factor merely rescales equilibrium abundances, leaving their relative proportions and equilibrium diversity unchanged. A similar invasion criterion can be derived for a simplified two-species Law–Watkinson model [6]. When species *j* is at its monoculture equilibrium, 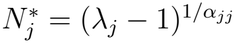, species *i* can increase when rare if

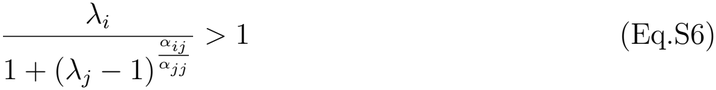

When 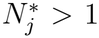 (equivalently, *λ*_*j*_ *>* 2), reducing *α*_*ij*_ makes invasion easier, all else being equal, and thus increases the likelihood of coexistence. We further tested this by examining the diversity consequences of generally weaker heterospecific competition in our fitted seven-species Law–Watkinson model, which should promote coexistence in multispecies systems [11]. We applied the model-estimated dynamic parameters from our system’s control plots in a deterministic simulation, separately for each posterior draw. We then applied the same simulations, but multiplied all heterospecific competition parameters *α*_*ij*_ (*i*≠*j*) by 0.5, leaving all other parameters unchanged. The resulting equilibrium diversity is substantially higher where a uniform reduction was applied to all the heterospecific competition parameters (Fig. S1).

The main simplification in our rudimentary interpretation of the general competition hypothesis is assuming that disturbance reduces all heterospecific competition coefficients by the same proportion. When this is not the case, the consequences for diversity depend on which competitive interactions are weakened and by how much. For example, in communities with asymmetric competition, weakening the effects of strong competitors on weaker species may increase diversity, whereas weakening the effects of weaker competitors on strong competitors may have the opposite effect. The abundance of the species involved also matters, as reductions in the competitive effects of common species may have greater consequences than similar reductions involving rare species.

**Figure S1:**
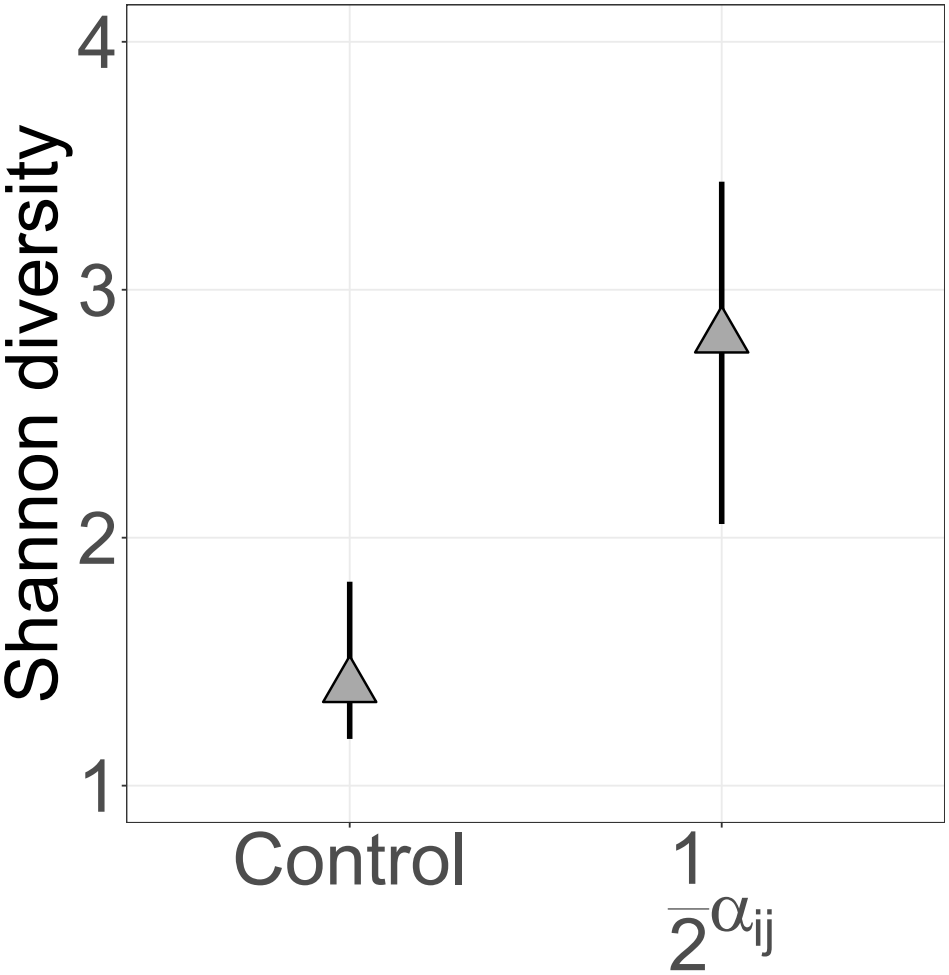
Reducing heterospecific competition parameters by a common factor increases equilibrium diversity. Triangles represent the posterior median, and bars represent the 90% credible interval. Control indicates where all heterospecific competition parameters are as estimated for the control plots by the model, and ^1^ *α*_*ij*_ represents where all these parameters are halved. The simulations were deterministic, starting from *N*_*i,t*_ = 10 and *N*_*i,t*−1_ = 5 for all species. The median change in abundance over the final four time steps was zero, consistent with convergence of the simulated community. Diversity is expressed as the effective number of species, exp(*H*^′^), where *H*^′^ is Shannon entropy. Median species diversity is doubled where all heterospecific competition parameters are halved, with non-overlapping 90% credible intervals.

#### Synthesis

The two pathways might not be mutually exclusive: in model formulations that are often used to describe natural populations, equilibrium abundance and invasion can depend on intrinsic growth rates and interactions (Eq.S6). Disturbance may simultaneously change population growth rates and the interaction matrix, and thus the diversity consequences of one pathway may depend on the other [12, 13]. Neither pathway’s individual effect necessarily predicts its contribution when both operate together. Furthermore even if only one pathway operates, demographic and competition parameters must be examined, since in reality the effects may not be uniform and depend on the model.

To test whether these mechanisms operate in natural communities, we quantified how disturbance modifies all demographic and competitive parameters within a dynamic community model [14]. Counterfactual simulations then allow us to selectively apply disturbance effects to particular parameters while keeping others at their control values, thereby isolating their contributions to diversity and distinguishing whether disturbance alters the proposed mechanisms from whether these changes actually increase diversity [15]. The main advantage of this approach is that it incorporates differential species responses, the number of competing species, and the structure of their interactions into the predictions, rather than “assuming away” these complications. The resulting predictions remain conditional on how well the chosen model captures the dynamics of the community.

### 2 Supplementary Methods

This appendix provides additional details on the experimental system (Section 2.1), model development and parameterization (Section 2.2), and equilibrium and counterfactual simulations (Section 2.3).

#### 2.1 Additional experimental details

The containers were filled with Hamra soil collected from deeper than approximately 1 m at a nearby site. Soil was collected from depth to minimize the introduction of seeds from the resident vegetation. Seeds of all seven experimental species were collected from nearby natural Hamra communities and sown only when the experiment was established.

Before sowing, germination trials were conducted in a phytotron under cool, short-day conditions representative of the local winter growing season. Species-specific germination fractions from these trials were used together with the target relative frequencies of each initial-composition treatment to determine the number of seeds sown. We aimed for a total initial density of approximately 3×10^3^ seedlings m^−2^. Realized seedling densities differed from these targets at the first growing season, when exceptionally heavy rainfall visibly displaced seeds and reduced establishment of several species. Densities increased substantially during the second growing season, reaching several thousand individuals per square meter.

The experimental communities grew outdoors and were therefore exposed to natural seasonal and interannual climatic variation. Because soil in containers dries more rapidly than continuous field soil, supplementary irrigation was applied equally to all communities during prolonged dry periods, with the amount adjusted according to conditions to prevent widespread drought-induced mortality. The soil collected from depth also contained little organic matter and was relatively nutrient poor. To prevent strong nutrient limitation from dominating the competitive dynamics, all containers received slow-release NPK fertilizer with micronutrients (Osmocote Plus, 15-9-12 N-P_2_O_5_-K_2_O) at the beginning of each growing season, corresponding to approximately 5 g N m^−2^.

Containers were separated by either approximately 1 or 2.2 m, with no systematic association between spacing and experimental treatment. Vegetation growing between containers was regularly removed by manual weeding, mowing, or herbicide application to reduce weeds and external seed input. Nets surrounding the containers were closed toward the end of the growing season to minimize seed movement among communities and reopened following the first germinating rains of the subsequent season.

The permanent 40 cm × 40 cm census quadrat was subdivided into four 20 cm × 20 cm sections to facilitate counting. In the second experimental year, a subset of these sections could not be sampled because of limited personnel availability. In affected *B. scoparius* monocultures and mixture plots, *B. scoparius* was counted in two of the four sections; in the same affected mixture plots, *H. spontaneum* was also counted in only two sections. For these observations, counts from the two sampled sections were summed and multiplied by two to estimate abundance for the full 40 cm × 40 cm quadrat.

Plot 70 was not sampled in the second experimental year and was therefore excluded from the analysis. Because year 2 enters all three consecutive-year sets used for model fitting, as (*t* + 1) in the first transition, (*t*) in the second, and (*t* − 1) in the third, its absence prevented construction of any complete transition sequence for this plot. In addition, two *L. halophilus* monocultures, plots 144 and 182, failed to germinate during the first year. These plots were excluded from the first transition because their observed abundance at (*t* − 1) was zero and Eq. (2) therefore deterministically predicts zero abundance at (*t* + 1).

#### 2.2 Model fitting and diagnostics

The model was estimated using three Markov chains where `adapt_delta = 0.95`, each run for 2 × 10^4^ steps including 2 × 10^3^ warm-up iterations. Every tenth post-warm-up draw was retained, thus keeping 5.4 × 10^3^ posterior draws. Convergence was validated by ensuring that Bulk & Tail Effective Sample Size (ESS) ≥ 4 × 10^2^ and 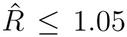for all parameters [16]. Across all parameters the maximum 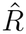 was 1.001936, while the minimum bulk ESS and minimum tail ESS both exceeded 4 × 10^3^, indicating excellent overall convergence. No divergent transitions occurred, and no iterations reached the maximum tree depth. For most parameters, initial values were set at the centers of their prior distributions to avoid initializing the Markov chains in the extreme tails of the parameter space. Absolute goodness-of-fit was tested using the `DHARMa` package [17], and by simulating the model to project an observed time step using the previous ones (Fig. 2), a method which Stouffer [18] highlights as a valuable indicator of whether the model truly captures how the system operates.

#### 2.2.1 Model design and selection

We looked for the most parsimonious model that fits the experimental annual census data. Our criteria were for it to have biological meaning, to provide a good representation of our data, including the experimental regimes and year-to-year variability, and to be favorable in model selection. For the first purpose, we considered three classical functional forms: Law & Watkinson [6], Beverton & Holt [4], and Ricker [5]. For the second purpose, we included the experimental treatments’ effects on each parameter.

Additionally, we *a priori* knew that Mediterranean systems show high climatic temporal variability [19], and indeed the four experimental years varied greatly in precipitation, as well as average daily temperature highs and lows (Table S1). While supplementary irrigation was applied in dry years to sustain the experimental communities, differences in natural precipitation may alter other environmental factors, e.g., VPD and radiation, thereby affecting plant growth and responses to biotic and abiotic factors. To keep the model as simple as possible, we did not want to explicitly account for these differences; thus, we included a hierarchical year effect structure for all the relevant parameters (see SI 2.2.3 for details).

**Table S1:** Temperature and precipitation conditions vary between sampling years. Temperature values are daily means calculated from October 1 of the preceding calendar year through April 30 of the sampling year. Precipitation is the total recorded during the corresponding rain season, defined as August 1 of the preceding calendar year through July 31 of the sampling year. For example, sampling year 2025 corresponds to the 2024/25 rain season. Temperature and precipitation data were obtained from the Israel Meteorological Service. Both were recorded at the Bet Dagan station (station 136741; coordinates 32.0073^°^N, 34.8138^°^E, 11.3 km aerial distance from the experiment; elevation 31 m). Mean rain-season precipitation at the station is 539.3 mm.

| Sampling year | Precipitation (mm) | Mean high (°C) | Mean low (°C) |
|---|---|---|---|
| 2022 | 780.1 | 22.5 | 11.0 |
| 2023 | 462.2 | 23.9 | 12.2 |
| 2024 | 604.0 | 24.5 | 13.3 |
| 2025 | 339.7 | 23.7 | 11.5 |

As model fit was not optimal, we tested two modifications to the regression: 1) the contribution of seeds from *t* − 1 (lagged response), representing the seed bank contribution as well as edge → center immigration (*s*_*i*_, Eq. 2), which incorporates the population size from *t* − 1 in addition to population size at *t*; and 2) a different noise parameterization from the classical negative binomial. The introduction of a new noise parameterization stemmed from strange model behavior around small *N*_*t*−1_ values, implying high uncertainty around them. The new noise parameterization is detailed in SI 2.2.2.

We compared all three functional forms under both noise parameterizations and with or without the inclusion of the memory parameter using `loo_compare` from the loo package [20]. The three baseline models, without either addition, exhibited substantial sampling and convergence issues and were therefore excluded from the LOO comparison. The Law-Watkinson model with the seed-bank addition and the new noise parameterization was the best-ranking form [21] (Table S2).

Aldebert & Stouffer [22] stress that different functional forms could qualitatively alter the predictions of coexistence or exclusions, and that different species may be better described by different functional forms. This highlights one must be cautious in using a single functional form for a multispecies competition model. To examine whether our seven-species model may exhibit a good fit on average while poorly representing some species, we additionally fit a stripped-down version of the full model to each of the species, for the three functional forms (with hierarchical structures removed, as these can only be applied across species), examining each functional form’s predictive accuracy on the single-species level. For all species, Law-Watkinson is the highest-ranked form or statistically indifferent from that, while Beverton-Holt and Ricker and a significantly worse fit for some species (Table S3). Since Law-Watkinson is also the best fit across the system as a whole (Table S2), we selected this functional form.

**Table S2:** ∆ELPD comparison of the candidate models. Models were fit to all seven species simultaneously to estimate all parameters in the system, and include hierarchical heterospecific and year-to-year variation structures. Models are ordered from highest to lowest expected log predictive density (ELPD). ∆ELPD values are calculated relative to the highest-ranked model; a substantial difference is considered where |∆ELPD| *>* 4 and large relative to their associated standard error [21]. NP denotes the new noise parameterization, and SB denotes the seed-bank (lagged response) inclusion. The Law-Watkinson model under the new noise parameterization with the lagged response addition outperforms all other candidates.

| Model | ΔELPD | SE of ΔELPD |
|---|---|---|
| Law-Watkinson + NP + SB | 0.0 | 0.0 |
| Beverton-Holt + NP + SB | -50.9 | 10.5 |
| Ricker + NP + SB | -96.8 | 14.6 |
| Law-Watkinson + NP | -3163.9 | 247.5 |
| Beverton-Holt + NP | -3190.9 | 247.6 |
| Ricker + NP | -3290.3 | 247.9 |
| Law-Watkinson + SB | -5031.6 | 932.0 |
| Beverton-Holt + SB | -5066.6 | 927.8 |
| Ricker + SB | -5105.0 | 924.8 |

**Table S3:** ∆ELPD stripped-down model comparison for each species. The model was run to assess relative goodness-of-fit for each species separately (rather than assuming all species have the same functional form as in Table S2), *sans* any hierarchical structure (which can only be applied across species). Values are ∆ELPD + SE_∆_ relative to the best-supported model for that species; a substantial difference is considered where |∆ELPD| *>* 4 and large relative to their associated standard error [21]. For all species, Law-Watkinson is either the highest-ranked model or statistically indifferent from it, while Beverton-Holt is a substantially worse fit for *Hordeum* and *Trifolium* and Ricker is a substantially worse fit for *Bromus, Lotus*, and *Rumex*.

| Species | Law–Watkinson | Beverton–Holt | Ricker |
|---|---|---|---|
| Bromus | 0.0 + 0.0 | −7.6 + 5.2 | −45.6 + 11.3 |
| Hordeum | −5.9 + 9.0 | −11.0 + 9.1 | 0.0 + 0.0 |
| Lotus | 0.0 + 0.0 | −0.3 + 2.8 | −11.3 + 5.2 |
| Trifolium | 0.0 + 0.0 | −7.3 + 4.6 | −3.2 + 6.1 |
| Rumex | −0.9 + 4.3 | 0.0 + 0.0 | −31.7 + 7.6 |
| Ormenis | −2.5 + 4.9 | −0.4 + 5.6 | 0.0 + 0.0 |
| Silene | 0.0 + 0.0 | −0.7 + 4.7 | −1.5 + 4.9 |

##### 2.2.2 Noise parameterization

We ran the model under a negative binomial (NB) likelihood, as we handled noisy count data. This likelihood treats the variance as:

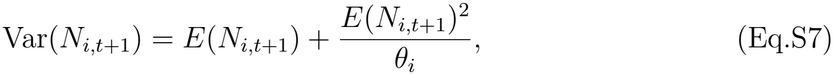

where *E*(*N*_*i,t*+1_) is the expected population average abundance for species *i* and *θ*_*i*_ is the inverse overdispersion parameter. However, this form introduced strange functional behavior near smaller *N*_*i,t*+1_ values. During model development we also applied a Quasi-Poisson (QP) variance structure [23] (*V ar*(*N*_*i,t*+1_) = *L*_*i*_ × *E*(*N*_*i,t*+1_)), which improved the fit for some of the species but impaired it for others. Consequently we used a different parameterization, where

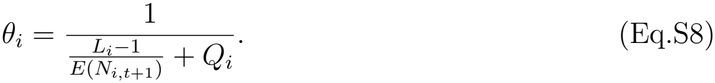

Substituting this back into the negative binomial variance, we get:

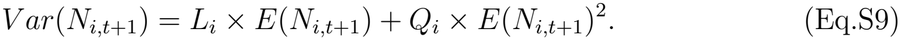

Where *L*_*i*_ and *Q*_*i*_ are scaling parameters for species *i* that can be considered the magnitude of demographic and environmental variability, respectively. This corresponds to the general agreement of the linear and quadratic components of stochasticity in ecological populations [24]. This parameterization will converge to NB when *L*_*i*_ = 1, and converge to QP when *Q*_*i*_ = 0. It was the most robust in model selection when compared to NB and was thus eventually used for the final model (Table S2).

##### 2.2.3 Hierarchical model structure

Each dynamic parameter was decomposed into three components: a baseline value, a multiplicative clipping effect, and a multiplicative year effect. For the intrinsic demographic parameters and conspecific competition, these components were species specific:

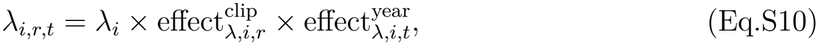

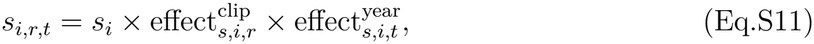

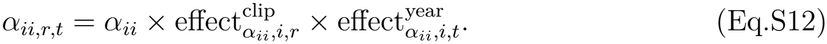

For heterospecific competition,

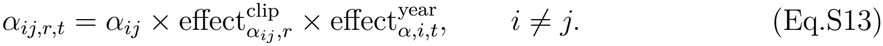

Here, *i* denotes the focal species, *j* the competing species, *r* the clipping regime, and *t* the annual transition. The baseline parameter describes the expected dynamics in the absence of clipping, while the two multiplicative terms describe deviations associated with clipping regime and year. The clipping effect was fixed to 1 for control communities, so values below or above 1 represent decreases or increases relative to the control, respectively.

Baseline heterospecific interaction coefficients were partially pooled across species pairs. Specifically,

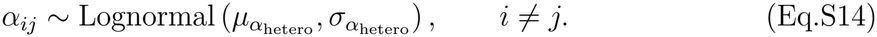

This hierarchical structure regularizes the 42 heterospecific interaction coefficients toward a common distribution while still allowing individual species pairs to differ when supported by the data.

Clipping effects on *λ*_*i*_, *s*_*i*_, and *α*_*ii*_ were estimated independently for each species and clipping regime:

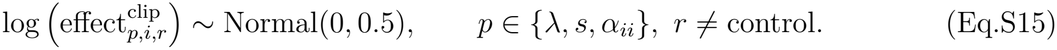

The treatment effects were exponentiated before entering the model as multiplicative effects. This prior is centered on no treatment effect, corresponding to a multiplicative effect of 1, and allows proportional increases and decreases symmetrically on the log scale. On the arithmetic scale, approximately 95% of the prior mass lies between 0.38 and 2.66, corresponding roughly to a three-fold decrease or increase in the parameter.

Because the 42 heterospecific interaction coefficients were themselves partially pooled, their clipping effects were also estimated hierarchically. For each clipping regime,

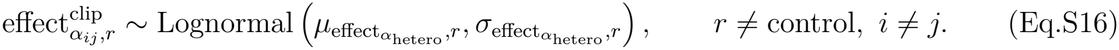

The hyperparameter 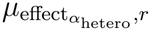 describes the overall effect of clipping regime *r* on heterospecific competition, whereas 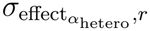 describes variation among individual species-pair responses. This structure allows us to estimate both a community-wide tendency for clipping to strengthen or weaken heterospecific competition and departures of individual interactions from that tendency.

The four experimental years differed substantially in precipitation and temperature (Table S1), so year effects were included to separate treatment responses from interannual environmental variation. For each parameter and species, the multiplicative year effect combined a component shared across species with an additional species-specific component:

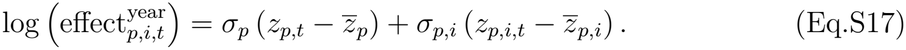

Here, *σ*_*p*_ determines the magnitude of interannual variation shared across species for parameter *p*, while *σ*_*p,i*_ allows species *i* to deviate from this shared response. The latent year effects were centered across years, so their mean on the log scale was zero and the geometric mean multiplicative year effect was therefore constrained to 1. Year effects could thus capture interannual deviations without changing the baseline scale of each parameter across the experimental period.

For heterospecific competition, we assumed that interannual environmental variation primarily altered the overall competitive sensitivity of each focal species rather than independently modifying its response to each competitor. Accordingly, all six heterospecific coefficients affecting the same focal species *i* shared the same species-specific year effect across competitors *j*. This reduced the number of year-specific interaction parameters while still allowing species to differ in how their competitive sensitivity changed among years. Priors for all baseline parameters, clipping effects, and year-effect hyperparameters are given in Section 2.2.4.

##### 2.2.4 Priors

*λ*_*i*_, *s*_*i*_, and *α*_*ii*_ were sampled on a log scale and exponentiated to ensure positivity. The priors were set to be:

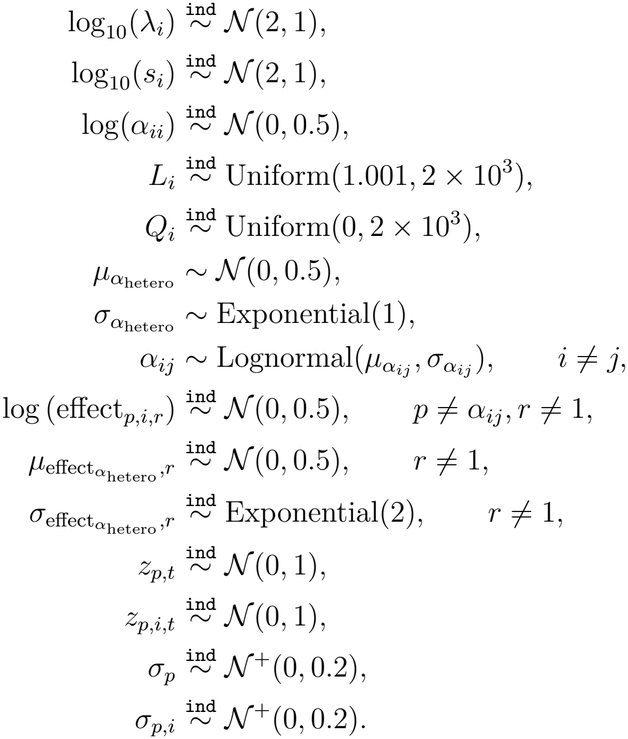

Italic subscripts are *i,j* = species, *p* = parameter, *r* = clipping regime, and *t* = year; ind stands for independently sampled vis-’a-vis the subscript index. For example, log_10_(*λ*_*i*_) is independently sampled seven times, as it’s only indexed by *i* and there are seven experimental species; log(effect_*p,i,r*_) is independently sampled 42 times, as it’s indexed by (*p* = 3) *λ, s, α*_*ii*_ × (*i* = 7) species × (*r* = 2) clipping regimes, = 42.

We aimed for informative priors, relying on *a priori* biological and ecological knowledge [25, 26]. The intrinsic potential population growth rates *λ*&*s* were given a prior centered around 10^2^, representing a conservative potential maximum for annual herbaceous plants, which commonly produce hundreds to thousands of seeds per individual under favourable, low-competition conditions [27], but giving lower, equal probabilities to 10^1^ and 10^3^, allowing a wide-enough range for r or K strategists. All the interaction coefficients (*α*) were given a prior centered around 1 on the arithmetic scale, to assume compensating dynamics but allow for under- or over-compensation [28], and constrained to be positive, as in the Law-Watkinson model (Eq. 2) negative *α* values would lead to a biologically invalid conclusion, where a species’ negative effect on population growth decreases as its population size increases; still this model allows only competitive interactions, which prevail in these Mediterranean annual systems [29]. For the variance parameters *L&Q* we used non-informative priors as we had no informed knowledge of them, but *L* had a lower bound 1.001 to avoid a negative *θ* (Eq.S8) while still allowing approximation to a negative-binomial variance structure (2.2.2). Clipping effects were given priors centered on zero on the log scale, corresponding to a multiplicative effect of 1 after exponentiation, so that *a priori* the clipping regimes are assumed to have no effect while allowing either decreases or increases to arise from the data. 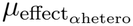 is likewise centered on zero on the log scale, corresponding to a multiplicative effect of 1 after exponentiation (Eq.S16). The year effects were modeled using a non-centred hierarchical parameterization. The latent annual deviations, *z*, were assigned standard-normal priors, *z* ∼ *N* (0, 1), defining the relative pattern of variation among years on a common scale. Their influence on each parameter was determined by the corresponding scale parameter, *σ*, which was assigned a half-normal prior, *σ* ∼ *N* ^+^(0, 0.2). This prior regularized interannual variation toward zero, equivalent to a multiplicative year effect of 1, while allowing stronger temporal variation when supported by the data. On the log scale, *σ* = 0.2 corresponds to a one-standard-deviation multiplicative change of approximately exp(*±*0.2) = 0.82–1.22. Separate global scale parameters described temporal variation shared among species, whereas species-specific scale parameters allowed each species to depart from the shared global response. For *α*_*ij*_, the species-specific year effect was shared among the six interaction coefficients affecting the same focal species. The scale parameters governing interannual variation in *α*_*ij*_ were additionally constrained to be below 0.25 to exclude implausibly large annual changes in interaction exponents and avoid numerical instability arising from the abundance terms raised to these exponents.

#### 2.3 Simulations

**Table S4:** Numerically invalid trajectories across posterior community simulations. Failures are defined as posterior draws producing NA, NaN, or Inf values. Cells show the mean (across simulations) percentage of trajectories excluded per community for each switchboard simulation. Such failures occurred in less than 0.2% of draws across simulations.

| Simulation scenario | Control | Early clipping | Late clipping |
|---|---|---|---|
| All effects on | 0.004% | 0.064% | 0.116% |
| Demographic hypothesis | 0.004% | 0.000% | 0.003% |
| General competition hypothesis | 0.004% | 0.151% | 0.128% |
| Grass → forb competition | 0.004% | 0.166% | 0.135% |
| Grasses dynamics | 0.004% | 0.026% | 0.032% |
| Grass → forb competition + grasses dynamics | 0.004% | 0.189% | 0.111% |

##### 2.3.1 Differentiation of equilibrium diversity patterns

To assess how strongly the experimental regimes differed in equilibrium diversity, we compared their posterior distributions using paired posterior draws. For each draw, we calculated the difference between each clipping treatment and the control, as well as the difference between early and late clipping. Positive values therefore indicate higher diversity under the first regime in each contrast, whereas negative values indicate higher diversity under the second.

We then calculated the smaller proportion of posterior differences falling below or above zero. Distributions centered around zero indicate little differentiation between regimes, while distributions lying predominantly on one side of zero indicate consistent posterior support for a difference. Results for all simulations are shown in Fig. S4.

### 3 Supplementary results

**Figure S2:**
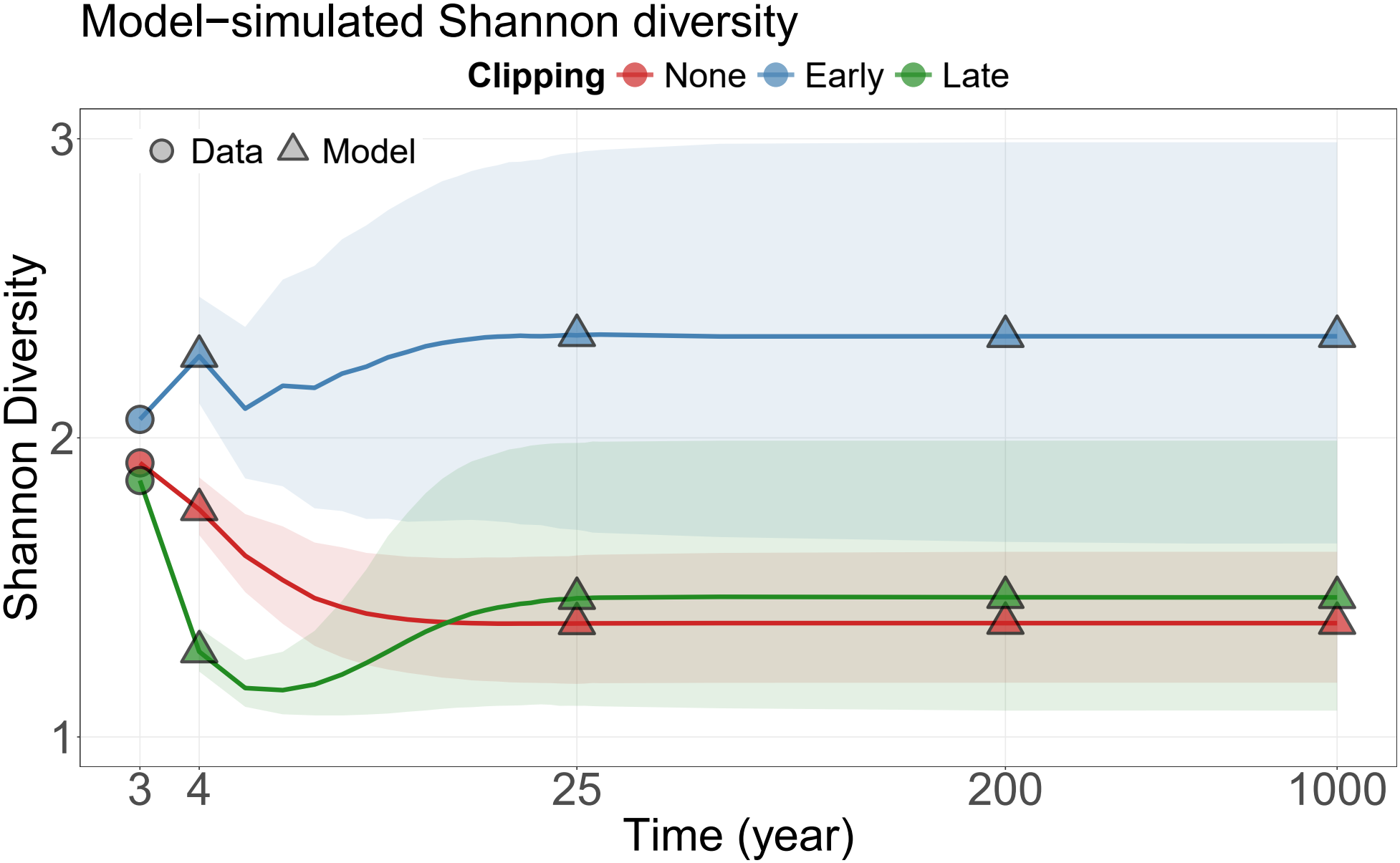
Equilibrium state is reached approximately by year 25. Diversity is expressed as the effective number of species, exp(*H*^′^), where *H*^′^ is Shannon entropy. Deterministic model projections initialized with observed population sizes from years 2 (*t* − 1) and 3 (*t*; circles), shown on a logarithmic time scale. Late clipping initially reduces diversity, followed by a partial recovery toward equilibrium. Triangles represent posterior medians at selected time steps, and shaded regions represent 90% credible intervals.

**Figure S3:**
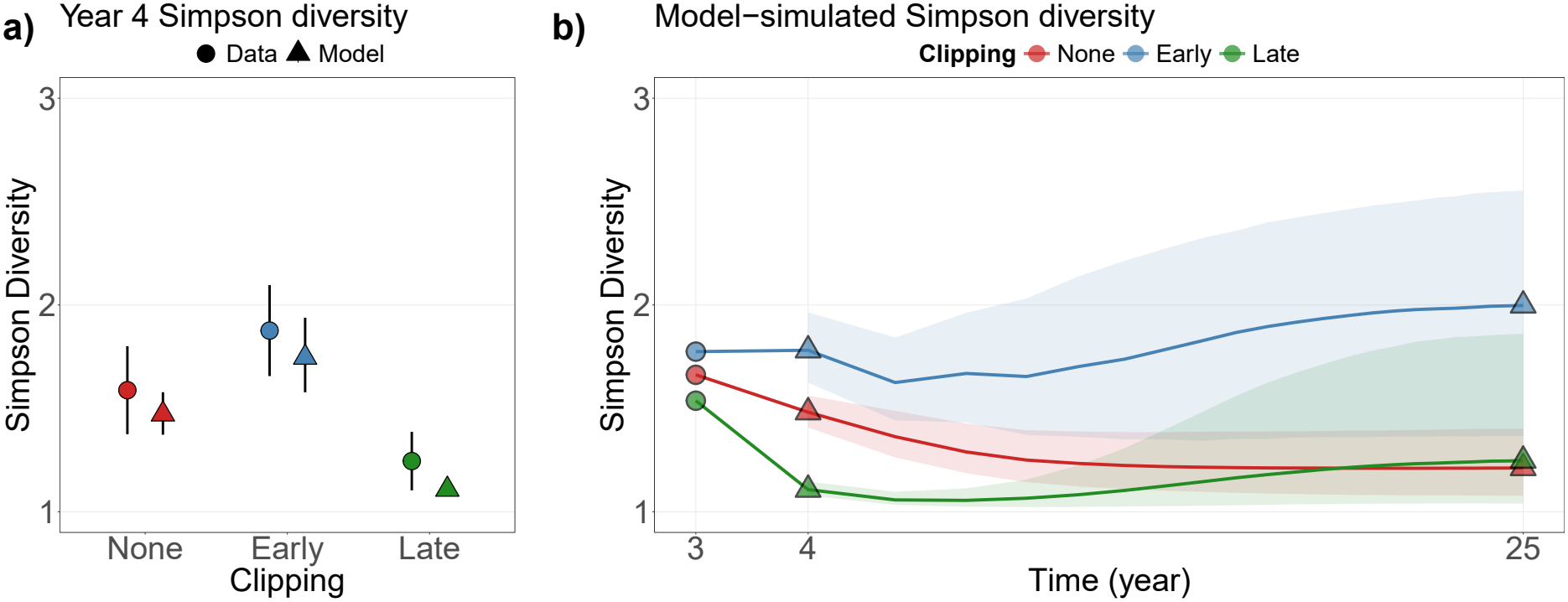
Early clipping shows higher Simpson diversity across timescales, while late clipping initially reduces diversity but this effect is diminished over time. Diversity was calculated as 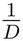, where *D* is Simpson concentration [30]. **a)** Observed year-4 diversity and corresponding in-sample posterior expectations. Points represent means, and error bars represent 90% confidence intervals for the data and 90% credible intervals for the model. **b)** Deterministic model projections initialized with observed population sizes from years 2 (*t* − 1) and 3 (*t*; circles), shown on a logarithmic time scale. Late clipping initially reduces diversity, followed by a partial recovery toward equilibrium. Triangles represent posterior medians at selected time steps, and shaded regions represent 90% credible intervals.

**Figure S4:**
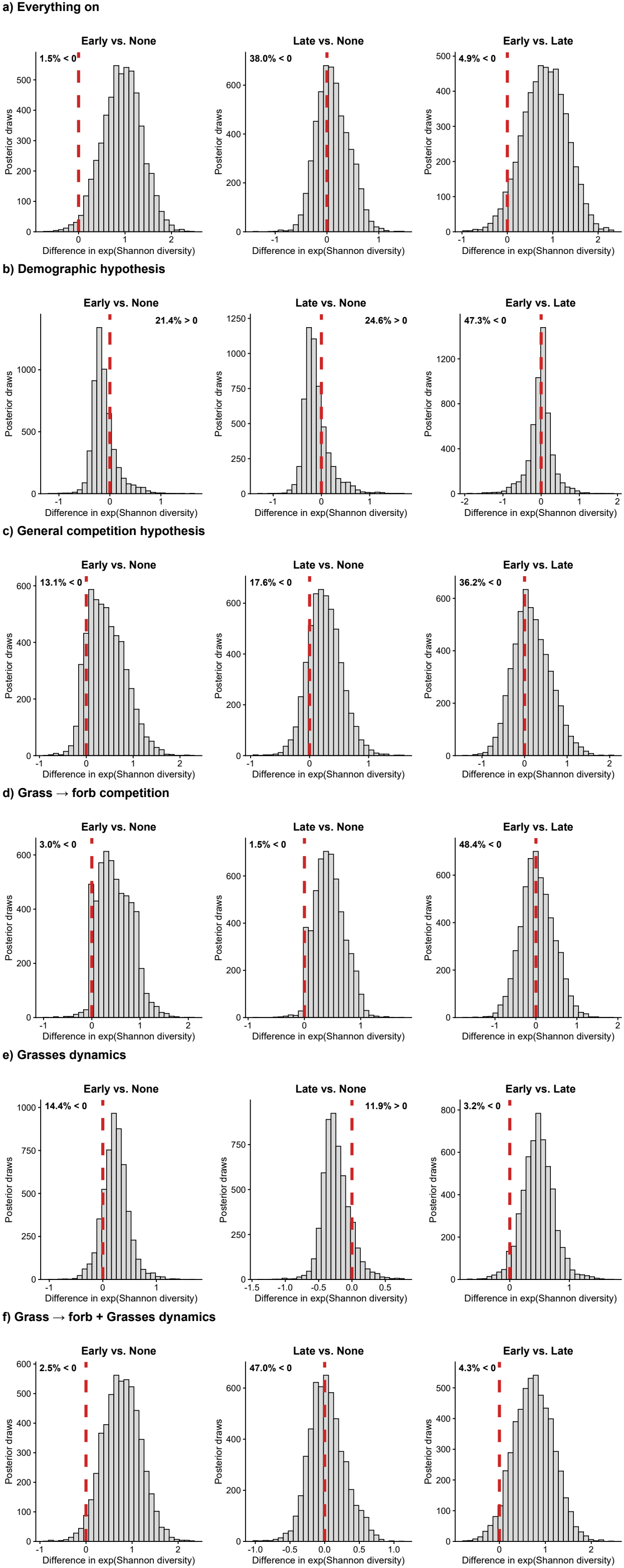
Posterior equilibrium diversity patterns show substantial differentiation among disturbance treatments across model configurations. Histograms show posterior distributions of paired, draw-level differences in exponentiated Shannon diversity at the final simulated timestep. Columns compare early clipping with no clipping, late clipping with no clipping, and early with late clipping, respectively. Positive values indicate greater diversity under the first treatment named in each comparison. Red dashed lines mark zero difference, and the percentage shown in each panel is the smaller posterior tail probability, reported as the proportion of differences below or above zero. Rows represent simulations where disturbance operates on **a)** all parameters, **b)** only the demographic rates *λ* &*s*, **c)** only heterospecific interactions *α*_*ij*_(*i*≠*j*), **d)** only grass (*j*) → forb (*i*) heterospecific interactions *α*_*ij*_, **e)** only grasses dynamics (*λ, s, α*_*ii*_&*α*_*ij*_(*i* ≠*j, i, j* = grass), and **f)** grass → forb interactions together with grasses dynamics.

**Figure S5:**
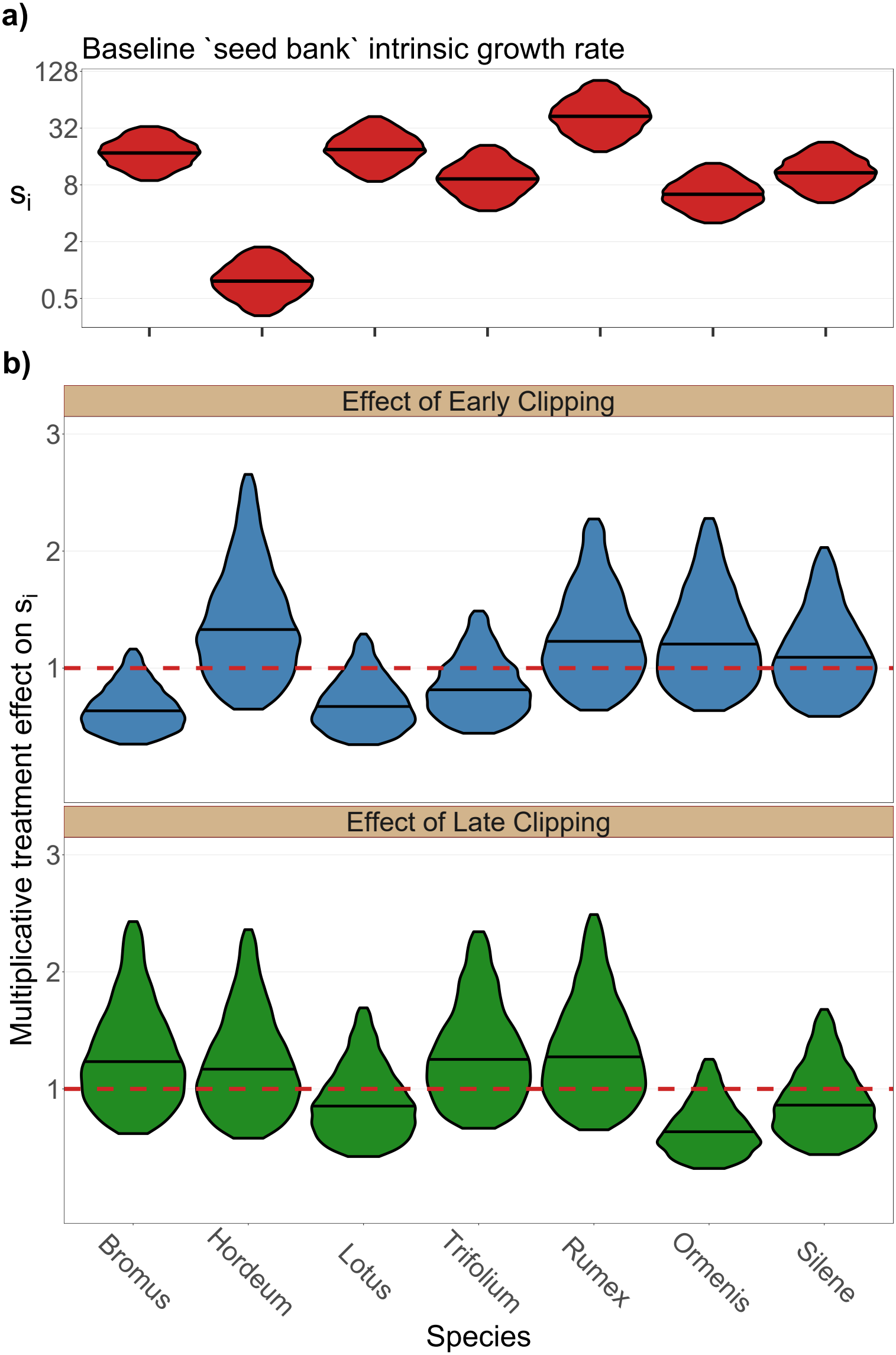
Disturbance does not decrease *s*_*i*_. Violins show the central 90% of posterior distributions; the line inside each violin is the median. Dashed red line in **b)** indicates no effect; effects above this line are positive, and below it are negative. **a)** Intrinsic seed bank rate *s*_*i*_ (i.e., the first term on the RHS in Eq.S11), for each of the seven experimental species in control plots on a logarithmic scale. **b)** Multiplicative effect of each disturbance regime on *s*_*i*_ (i.e., the second term on the RHS in Eq.S11). Disturbance does not generally reduce seed bank intrinsic growth rates; instead, the effect is inconsistent across species and treatments.

**Figure S6:**
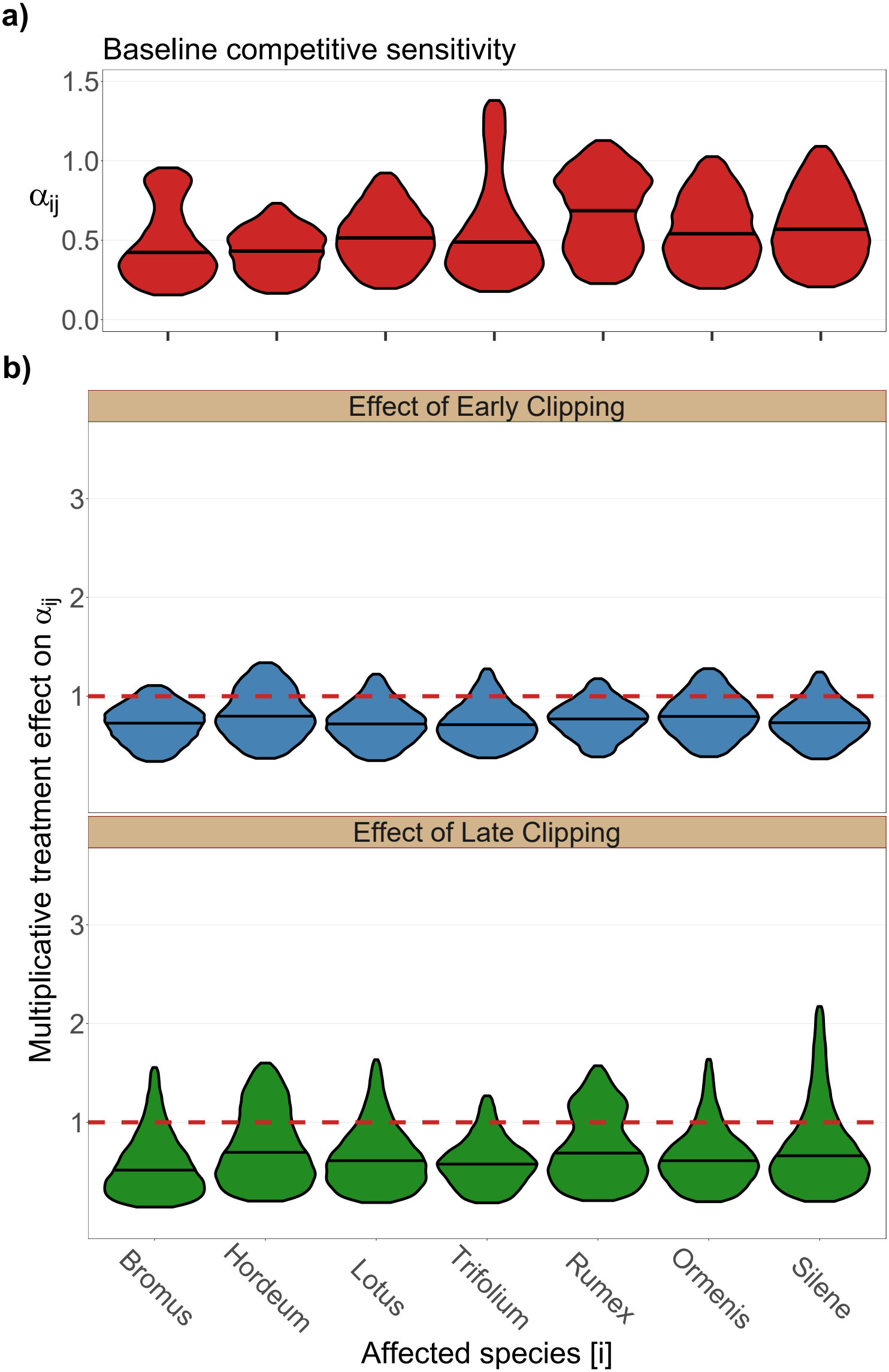
Disturbance reduces competitive sensitivity regardless of timing. Violins show the central 90% of posterior distributions; the line inside each violin is the median. **a)** Heterospecific competitive sensitivity *α*_*ij*_ (i.e., the first term on the RHS in Eq.S13), grouped by affected species *i* and pooled over all affecting species *j* ≠*i* in control plots. *Rumex* is the most sensitive competitor. **b)** shows the treatment effect (i.e., the second term on the RHS in Eq.S13). red lines in indicate no effect; effects above this line are positive, and below it are negative. Clipping generally weakens competitive sensitivity. In this figure the grouping of *α*_*ij*_ is done by affected species [j], while in Figure 4 of the main text it is done by affecting species i.

**Figure S7:**
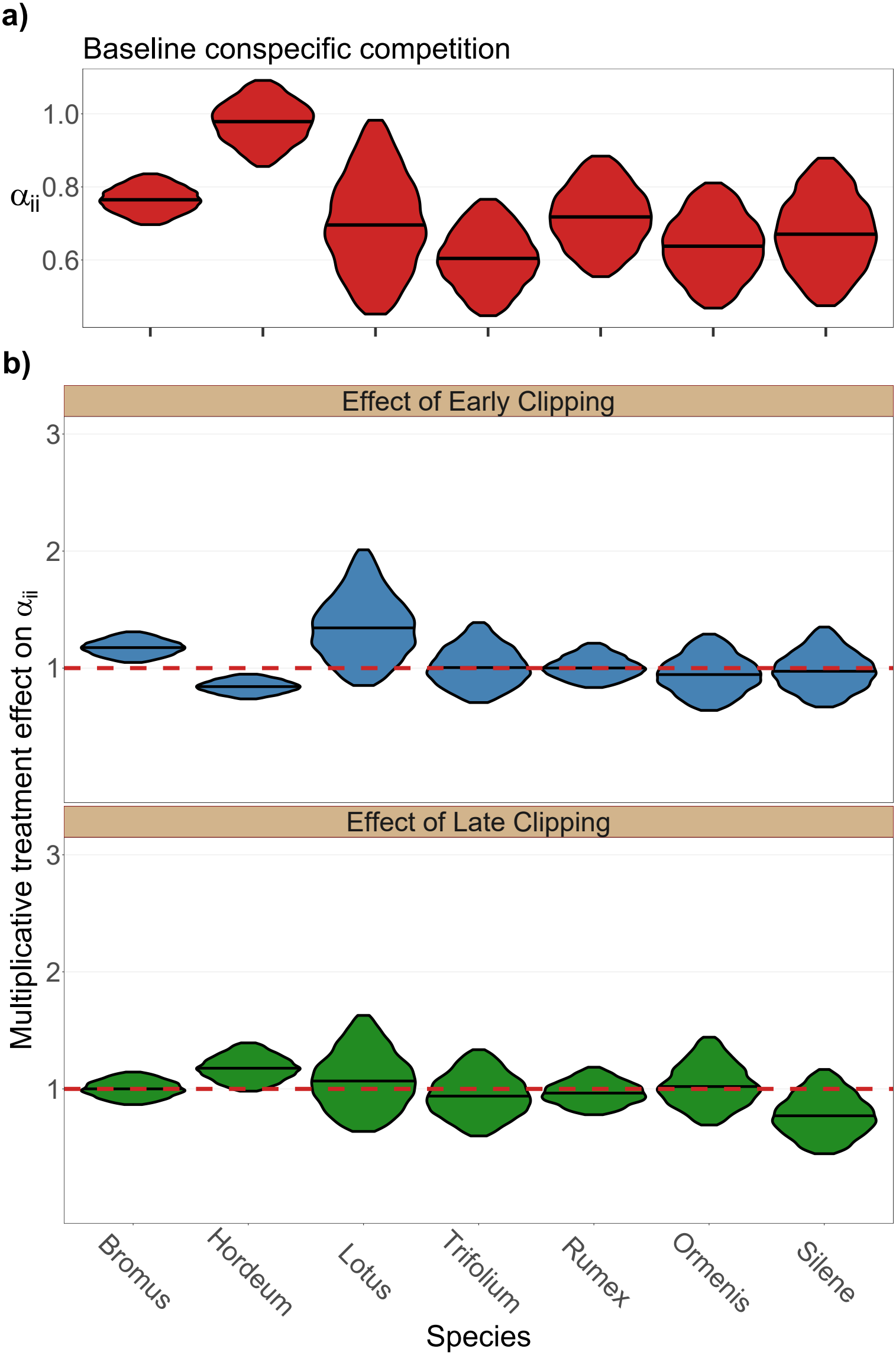
Disturbance does not decrease *α*_*ii*_. Violins show the central 90% of posterior distributions; the line inside each violin is the median. Dashed red line in **b)** indicates no effect; effects above this line are positive, and below it are negative. **a)** conspecific competition *α*_*ii*_ (i.e., the first term on the RHS in Eq.S12), for each of the seven experimental species in control plots. **b)** Multiplicative effect of each disturbance regime on *α*_*ii*_ (i.e., the second term on the RHS in Eq.S12). Disturbance does not generally reduce conspecific competition; instead, it’s mostly unchanged, but sometimes reduces or increases slightly.

**Figure S8:**
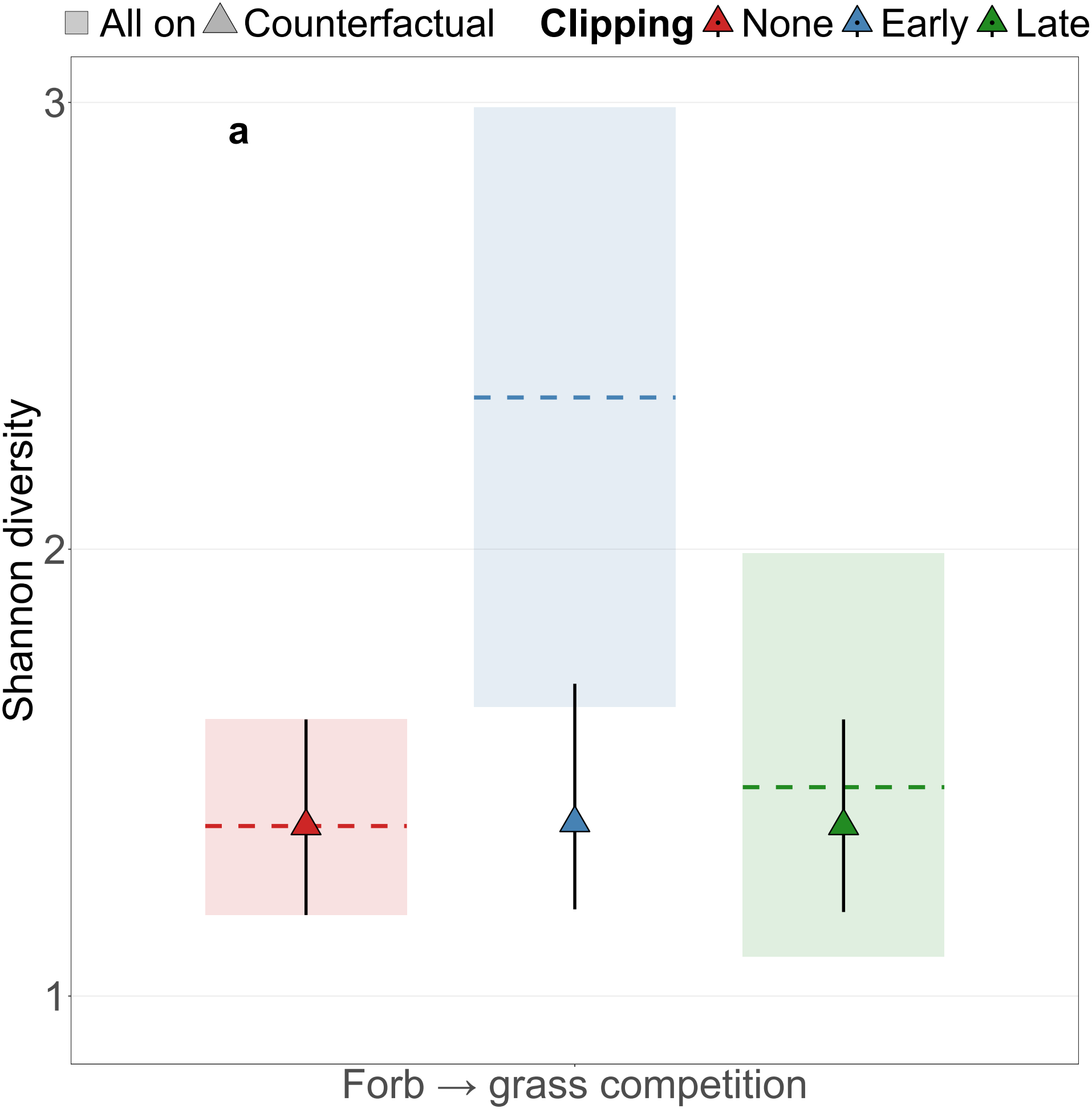
Counterfactual disturbances that only operate on the forb → forb, and forb → grasses competitive effects do not increase diversity. Simulations were run to equilibrium and initialized with observed abundances from years 2 (*t* − 1) and 3 (*t*). Rectangles and dashed lines represent the posterior medians and 90% credible intervals, respectively, of the full simulations, where disturbance acts on all the parameters. Triangles and error bars represent the posterior medians and 90% credible intervals, respectively, of the ‘what-if’ counterfactual simulations.

**Figure S9:**
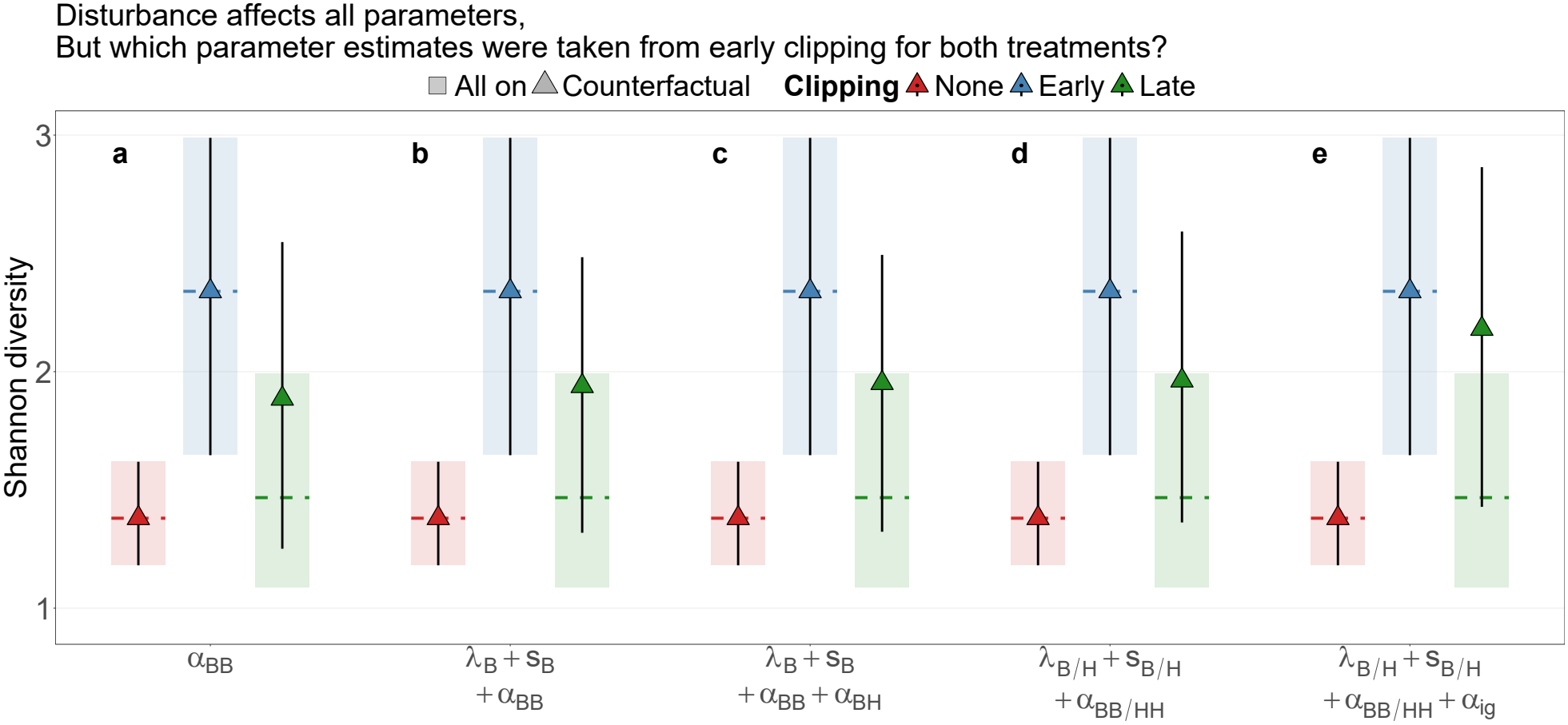
Increased diversity under early clipping compared to late clipping is almost entirely explained by the grasses’ intrinsic dynamics and competitive effects, but the intraspecific competition of *Bromus* is central in this. Simulations were run to equilibrium and initialized with observed abundances from years 2 (*t* − 1) and 3 (*t*). Rectangles and dashed lines represent the posterior medians and 90% credible intervals, respectively, of the full simulations, where disturbance acts on all the parameters and are completely identical in a-e. Triangles and error bars represent the posterior medians and 90% credible intervals, respectively, of the ‘what-if’ counterfactual simulations where subsets of the early clipping parameters were used for both early and late clipping. This helps pinpoint what parameters underlie the differences between these two treatments. **a)** If the conspecific competition of *Bromus* (Fig. S7) was the same under late clipping as it is under early clipping, late clipping diversity would increase. **b)** Pulling the population growth rates of *Bromus* (Fig. 4a,b, S5b) from early clipping for both treatments increases diversity even more, but slightly. **c)** Adding the competitive effect of *Hordeum* on *Bromus* to **b)** does not change late clipping diversity. **d)** Using all the grasses’ intrinsic dynamics from early clipping for both treatments increases late clipping diversity, but only marginally compared to **b). e)** If grasses’ intrinsic parameters and interactions and their competitive effects on each other and on the non-grass species would be affected by the late clipping as they are by the early clipping, diversity would increase almost identically under both treatments. This parameter subset is identical to the subset termed ‘Grass → forb + Grasses dynamics’ in Fig. 5e.

**Figure S10:**
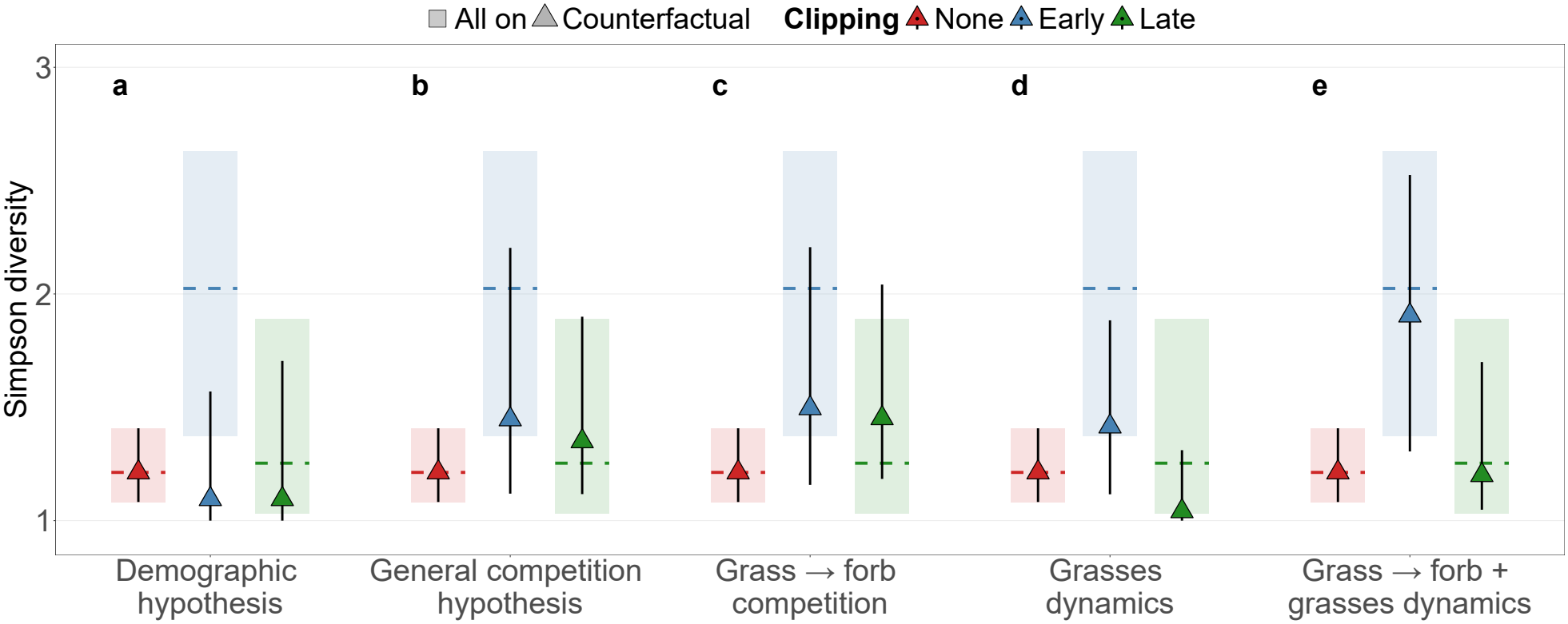
Increased equilibrium diversity under disturbance is explained by weakened grass → forb competition, while the overall response of diversity is explained by disturbance’s effect on the dynamics and effects of grass species on the community. Simulations were run to equilibrium and initialized with observed abundances from years 2 (*t* − 1) and 3 (*t*). Rectangles and dashed lines represent the posterior medians and 90% credible intervals, respectively, of the full simulations, where disturbance acts on all the parameters and are completely identical in a-e. Triangles and error bars represent the posterior medians and 90% credible intervals, respectively, of the ‘what-if’ counterfactual simulations. Diversity is expressed as the effective number of species,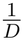, where *D* is Simpson concentration. **a)** If disturbance affected only demographic rates, equilibrium diversity would decrease under both clipping regimes. **b)** If disturbance affected only heterospecific competitive interactions, equilibrium diversity would increase under both clipping regimes. **c)** If disturbance affected only competitive interactions imposed by grasses on non-grass species, equilibrium diversity would increase under both clipping regimes. **d)** If disturbance affected only grasses’ intrinsic parameters and interactions, equilibrium diversity would show the same qualitative pattern as where disturbance acts through all parameters, but not the same quantitative pattern. **e)** If disturbance affected grasses’ intrinsic parameters and interactions and their competitive effect on the non-grass species, equilibrium diversity would show the same quantitative pattern as when disturbance acts through all parameters.

## Notes

### Competing Interest Statement

The authors have declared no competing interest.

### Summary of Updates

Abstract and discussion updated; Box 1 and SI 1 added.

## References

1. Ellis, E. C. et al. People have shaped most of terrestrial nature for at least 12,000 years. Proceedings of the National Academy of Sciences 118, e2023483118 (2021).

2. Stephens, L. et al. Archaeological assessment reveals Earth’s early transformation through land use. Science 365, 897–902 (2019).

3. Gossner, M. M. et al. Land-use intensification causes multitrophic homogenization of grassland communities. Nature 540, 266–269 (2016).

4. Foley, J. A. et al. Global Consequences of Land Use. Science 309, 570–574 (2005).

5. Tilman, D. & Lehman, C. Human-caused environmental change: Impacts on plant diversity and evolution. Proceedings of the National Academy of Sciences 98, 5433–5440 (2001).

6. Grime, J. P. Competitive Exclusion in Herbaceous Vegetation. Nature 242, 344–347 (1973).

7. Huston, M. A. A General Hypothesis of Species Diversity. The American Naturalist 113 (1979).

8. Connell, J. H. Diversity in Tropical Rain Forests and Coral Reefs. Science 199 (1978).

9. Hillebrand, H. et al. Consumer versus resource control of producer diversity depends on ecosystem type and producer community structure. Proceedings of the National Academy of Sciences 104, 10904–10909 (2007).

10. Worm, B., Lotze, H. K., Hillebrand, H. & Sommer, U. Consumer versus resource control of species diversity and ecosystem functioning. Nature 417, 848–851 (2002).

11. Borer, E. T. et al. Herbivores and nutrients control grassland plant diversity via light limitation. Nature 508, 517–520 (2014).

12. Buisson, E., Archibald, S., Fidelis, A. & Suding, K. N. Ancient grasslands guide ambitious goals in grassland restoration. Science 377, 594–598 (2022).

13. Eskelinen, A., Harpole, W. S., Jessen, M.-T., Virtanen, R. & Hautier, Y. Light competition drives herbivore and nutrient effects on plant diversity. Nature 611, 301–305 (2022).

14. Fox, J. W. The intermediate disturbance hypothesis should be abandoned. Trends in Ecology & Evolution 28, 86–92 (2013).

15. Huston, M. A. Disturbance, productivity, and species diversity: empiricism vs. logic in ecological theory. Ecology 95, 2382–2396 (2014).

16. Kondoh, M. Unifying the relationships of species richness to productivity and disturbance. Proceedings of the Royal Society B: Biological Sciences 268, 269–271 (2001).

17. Kadmon, R. & Benjamini, Y. Effects of Productivity and Disturbance on Species Richness: A Neutral Model. The American Naturalist 167 (2006).

18. Belsky, A. J. Effects of grazing, competition, disturbance and fire on species composition and diversity in grassland communities. Journal of Vegetation Science 3, 187–200 (1992).

19. Koerner, S. E. et al. Change in dominance determines herbivore effects on plant biodiversity. Nature Ecology & Evolution 2, 1925–1932 (2018).

20. Klaus, V. H. et al. Enriching plant diversity in grasslands by large-scale experimental sward disturbance and seed addition along gradients of land-use intensity. Journal of Plant Ecology 10, 581–591 (2017).

21. Sternberg, M. et al. Testing the limits of resistance: a 19-year study of Mediterranean grassland response to grazing regimes. Global Change Biology 21, 1939–1950 (2015).

22. Gross, K. L., Mittelbach, G. G. & Reynolds, H. L. Grassland invasibility and diversity: Responses to nutrients, seed input, and disturbance. Ecology 86, 476–486 (2005).

23. Proulx, M. & Mazumder, A. Reversal of Grazing Impact on Plant Species Richness in Nutrient-Poor Vs. Nutrient-Rich Ecosystems. Ecology 79, 2581–2592 (1998).

24. Shea, K., Roxburgh, S. H. & Rauschert, E. S. J. Moving from pattern to process: coexistence mechanisms under intermediate disturbance regimes. Ecology Letters 7, 491–508 (2004).

25. Volterra, V. Variazioni e fluttuazioni del numero d’individui in specie animali conviventi 148 pp. (Societá anonima tipografica “Leonardo da Vinci”, 1927).

26. Novak, M. et al. Characterizing Species Interactions to Understand Press Perturbations: What Is the Community Matrix? Annual Review of Ecology, Evolution, and Systematics 47, 409–432 (2016).

27. Stouffer, D. B. A critical examination of models of annual-plant population dynamics and density-dependent fecundity. Methods in Ecology and Evolution 13, 2516–2530 (2022).

28. Bråthen, K. A., Pugnaire, F. I. & Bardgett, R. D. The paradox of forbs in grasslands and the legacy of the mammoth steppe. Frontiers in Ecology and the Environment 19, 584–592 (2021).

29. DeMalach, N., Zaady, E. & Kadmon, R. Light asymmetry explains the effect of nutrient enrichment on grassland diversity. Ecology Letters 20, 60–69 (2017).

30. Nelson, R. A. et al. Forb diversity globally is harmed by nutrient enrichment but can be rescued by large mammalian herbivory. Communications Biology 8, 444 (2025).

31. Adler, P. B. et al. Competition and coexistence in plant communities: intraspecific competition is stronger than interspecific competition. Ecology Letters 21, 1319–1329 (2018).

32. Cervantes-Loreto, A. et al. Environmental context, parameter sensitivity, and structural sensitivity impact predictions of annual-plant coexistence. Ecological Monographs 93, e1592 (2023).

33. Godoy, O., Gómez-Aparicio, L., Matías, L., Pérez-Ramos, I. M. & Allan, E. An excess of niche differences maximizes ecosystem functioning. Nature Communications 11, 1–10 (2020).

34. Granjel, R. R., Allan, E. & Godoy, O. Nitrogen enrichment and foliar fungal pathogens affect the mechanisms of multispecies plant coexistence. New Phytologist 237, 2332–2346 (2023).

35. Hess, C., Levine, J. M., Turcotte, M. M. & Hart, S. P. Phenotypic plasticity promotes species coexistence. Nature Ecology and Evolution 6, 1256–1261 (2022).

36. Napier, J. D., Mordecai, E. A. & Heckman, R. W. The role of drought- and disturbance- mediated competition in shaping community responses to varied environments. Oecologia 181, 621–632 (2016).

37. Terry, J. C. D. & Armitage, D. W. Widespread analytical pitfalls in empirical coexistence studies and a checklist for improving their statistical robustness (2024).

38. Levine, J. M. & Hillerislambers, J. The importance of niches for the maintenance of species diversity. Nature 461 (2009).

39. Kraft, N. J., Godoy, O. & Levine, J. M. Plant functional traits and the multidimensional nature of species coexistence. Proceedings of the National Academy of Sciences of the United States of America 112, 797–802 (2015).

40. Tredennick, A. T., Hooten, M. B. & Adler, P. B. Do we need demographic data to forecast plant population dynamics? Methods in Ecology and Evolution 8, 541–551 (2017).

41. Facelli, J. M. & Pickett, S. T. A. Plant litterits dynamics and effects on plant community structure. Botanical Review 57, 1–32 (1991).

42. Jessen, M.-T., Auge, H., Harpole, W. S. & Eskelinen, A. Litter accumulation, not light limitation, drives early plant recruitment. Journal of Ecology 111, 1174–1187 (2023).

43. Newman, E. I. Competition and Diversity in Herbaceous Vegetation. Nature 244, 310– 310 (1973).

44. Gurevitch, J. & Unnasch, R. S. Experimental removal of a dominant species at two levels of soil fertility. Canadian Journal of Botany 67, 3470–3477 (1989).

45. Lepš, J. Scale- and time-dependent effects of fertilization, mowing and dominant removal on a grassland community during a 15-year experiment. Journal of Applied Ecology 51, 978–987 (2014).

46. Segre, H., DeMalach, N., Henkin, Z. & Kadmon, R. Quantifying competitive exclusion and competitive release in ecological communities: A conceptual framework and a case study. PLoS ONE 11 (2016).

47. Zhang, P. et al. Dominant species predict plant richness and biomass in global grasslands. Nature Ecology & Evolution 9, 924–936 (2025).

48. Ellner, S. P., Snyder, R. E., Adler, P. B. & Hooker, G. An expanded modern coexistence theory for empirical applications. Ecology Letters 22, 3–18 (2019).

49. Adler, P. B. et al. What Have We Learned From Empirical Applications of Modern Coexistence Theory? Ecology Letters 29, e70404 (2026).

50. Issaka, D. S., Gross, O., Ayilara, I., Schabes, T. & DeMalach, N. Density-dependent and independent mechanisms jointly reduce species performance under nitrogen enrichment. Oikos 2023 (2023).

51. R Core Team. R: A Language and Environment for Statistical Computing R Foundation for Statistical Computing (Vienna, Austria, 2026).

52. Stan Development Team. RStan: the R interface to Stan 2025.

53. Hartig, F. DHARMa: Residual Diagnostics for Hierarchical (Multi-Level / Mixed) Regression Models (2026).

54. Vehtari, A. et al. loo: Efficient leave-one-out cross-validation and WAIC for Bayesian models 2025.

55. Wickham, H. et al. Welcome to the tidyverse. Journal of Open Source Software 4, 1686 (2019).

56. Law, R. & Watkinson, A. R. Response-Surface Analysis of Two-Species Competition: An Experiment on Phleum Arenarium and Vulpia Fasciculata. Journal of Ecology 75, 871–886 (1987).

57. DeMalach, N., Kigel, J. & Sternberg, M. The soil seed bank can buffer long-term compositional changes in annual plant communities. Journal of Ecology 109, 1275–1283 (2021).

58. Eskelinen, A. et al. Herbivory and nutrients shape grassland soil seed banks. Nature Communications 14, 3949 (2023).

59. Sivula, T., Magnusson, M., Matamoros, A. A. & Vehtari, A. Uncertainty in Bayesian leave-one-out cross-validation based model comparison. Bayesian Analysis (2025).

60. Beverton, R. J. H. & Holt, S. J. On the Dynamics of Exploited Fish Populations 541 pp. (Springer Science & Business Media, 1957).

61. Ricker, W. E. Effects of Compensatory Mortality upon Population Abundance. The Journal of Wildlife Management The Journal of Wildlife Management, 45–51 (1954).

62. Giorgi, F. & Lionello, P. Climate change projections for the Mediterranean region. Global and Planetary Change. Mediterranean climate: trends, variability and change 63, 90–104 (2008).

63. Jost, L. Entropy and diversity. Oikos 113, 363–375 (2006).

## References

1. Grime, J. P. Competitive Exclusion in Herbaceous Vegetation. Nature 242, 344–347 (1973).

2. Huston, M. A. A General Hypothesis of Species Diversity. The American Naturalist 113 (1979).

3. Volterra, V. Variazioni e fluttuazioni del numero d’individui in specie animali conviventi 148 pp. (Societá anonima tipografica “Leonardo da Vinci”, 1927).

4. Beverton, R. J. H. & Holt, S. J. On the Dynamics of Exploited Fish Populations 541 pp. (Springer Science & Business Media, 1957).

5. Ricker, W. E. Stock and Recruitment. Journal of the Fisheries Research Board of Canada 11, 559–623 (1954).

6. Law, R. & Watkinson, A. R. Response-Surface Analysis of Two-Species Competition: An Experiment on Phleum Arenarium and Vulpia Fasciculata. Journal of Ecology 75, 871–886 (1987).

7. Kondoh, M. Unifying the relationships of species richness to productivity and disturbance. Proceedings of the Royal Society B: Biological Sciences 268, 269–271 (2001).

8. Kadmon, R. & Benjamini, Y. Effects of Productivity and Disturbance on Species Richness: A Neutral Model. The American Naturalist 167 (2006).

9. Arditi, R., Tyutyunov, Y. V., Titova, L. I., Rohr, R. P. & Bersier, L.-F. The Dimensions and Units of the Population Interaction Coefficients. Frontiers in Ecology and Evolution 9 (2021).

10. Broekman, M. J. E. et al. Signs of stabilisation and stable coexistence. Ecology Letters 22, 1957–1975 (2019).

11. Kessler, D. A. & Shnerb, N. M. Generalized model of island biodiversity. Physical Review E 91, 042705 (2015).

12. Saavedra, S. et al. A structural approach for understanding multispecies coexistence. Ecological Monographs 87, 470–486 (2017).

13. Löffler, T. J. & Lischke, H. Changing relative intrinsic growth rates of species alter the stability of species communities. Frontiers in Ecology and Evolution 11 (2023).

14. Hart, S. P., Freckleton, R. P. & Levine, J. M. How to quantify competitive ability. Journal of Ecology 106, 1902–1909 (2018).

15. Adler, P. B. et al. What Have We Learned From Empirical Applications of Modern Coexistence Theory? Ecology Letters 29, e70404 (2026).

16. Vehtari, A., Gelman, A., Simpson, D., Carpenter, B. & Bürkner, P.-C. Rank-Normalization, Folding, and Localization: An Improved R for Assessing Convergence of MCMC (with Discussion). Bayesian Analysis 16, 667–718 (2021).

17. Hartig, F. DHARMa: Residual Diagnostics for Hierarchical (Multi-Level / Mixed) Regression Models (2026).

18. Stouffer, D. B. A critical examination of models of annual-plant population dynamics and density-dependent fecundity. Methods in Ecology and Evolution 13, 2516–2530 (2022).

19. Giorgi, F. & Lionello, P. Climate change projections for the Mediterranean region. Global and Planetary Change. Mediterranean climate: trends, variability and change 63, 90–104 (2008).

20. Vehtari, A. et al. loo: Efficient leave-one-out cross-validation and WAIC for Bayesian models 2025.

21. Sivula, T., Magnusson, M., Matamoros, A. A. & Vehtari, A. Uncertainty in Bayesian leave-one-out cross-validation based model comparison. Bayesian Analysis (2025).

22. Aldebert, C. & Stouffer, D. B. Community dynamics and sensitivity to model structure: towards a probabilistic view of process-based model predictions. Journal of the Royal Society Interface 15, 20180741 (2018).

23. Ver Hoef, J. M. &Boveng, P. L. Quasi-Poisson Vs. Negative Binomial Regression: How Should We Model Overdispersed Count Data? Ecology 88, 2766–2772 (2007).

24. Kalyuzhny, M. et al. Temporal fluctuation scaling in populations and communities. Ecology 95, 1701–1709 (2014).

25. Banner, K. M., Irvine, K. M. & Rodhouse, T. J. The use of Bayesian priors in Ecology: The good, the bad and the not great. Methods in Ecology and Evolution 11, 882–889 (2020).

26. McElreath, R. Statistical Rethinking: A Bayesian Course with Examples in R and STAN (CRC Press LLC, Milton, UNITED KINGDOM, 2020).

27. Hodgson, J. G. et al. Seed size, number and strategies in annual plants: a comparative functional analysis and synthesis. Annals of Botany 126, 1109–1128 (2020).

28. Bagchi, R. et al. Testing the Janzen-Connell mechanism: pathogens cause overcompen-sating density dependence in a tropical tree. Ecology Letters 13, 1262–1269 (2010).

29. Issaka, D. S., Gross, O., Ayilara, I., Schabes, T. & DeMalach, N. Density-dependent and independent mechanisms jointly reduce species performance under nitrogen enrichment. Oikos 2023 (2023).

30. Jost, L. Entropy and diversity. Oikos 113, 363–375 (2006).

